# A pseudohelicase-centered complex couples assembly-dependent RNA cleavage to poly(UG)ylation

**DOI:** 10.64898/2026.09.04.749402

**Authors:** Virginia Busetto, Martin Brehm, Annamaria Sgromo, Maximilian Mager, Lizaveta Pshanichnaya, Elena Becker, René F. Ketting, Stefan Ameres, Sebastian Falk

## Abstract

Robust RNA interference in *Caenorhabditis elegans* requires Argonaute-targeted RNAs to be cleaved by RDE-8 and poly(UG)ylated by MUT-2, generating templates for RNA-dependent RNA polymerases that synthesize secondary small RNAs. RDE-8 and MUT-2 are physically coupled within the cleavage and poly(UG)ylation (CPUG) complex, together with the pseudonuclease NYN-1 and MUT-15, but how their activities are controlled and coordinated is unknown. Structural and biochemical analyses reveal that the RDE-8/NYN-1 nuclease module is autoinhibited, restricting cleavage outside CPUG. We identify MUT-15, the CPUG scaffold linking this nuclease module to MUT-2, as a pseudohelicase. MUT-15 remodels the nuclease module, licensing RDE-8 activity upon complex assembly. Reconstituted CPUG synthesizes long poly(UG) tails, with the substrate 3′ end determining tailing efficiency and register. Within CPUG, RNA engagement by MUT-2 limits RDE-8 access, while poly(UG) tails themselves are not cleaved by RDE-8, preventing re-cleavage. Together, CPUG couples assembly-dependent RNA cleavage to downstream processing, imposing safety and directionality.

**Graphical abstract:** 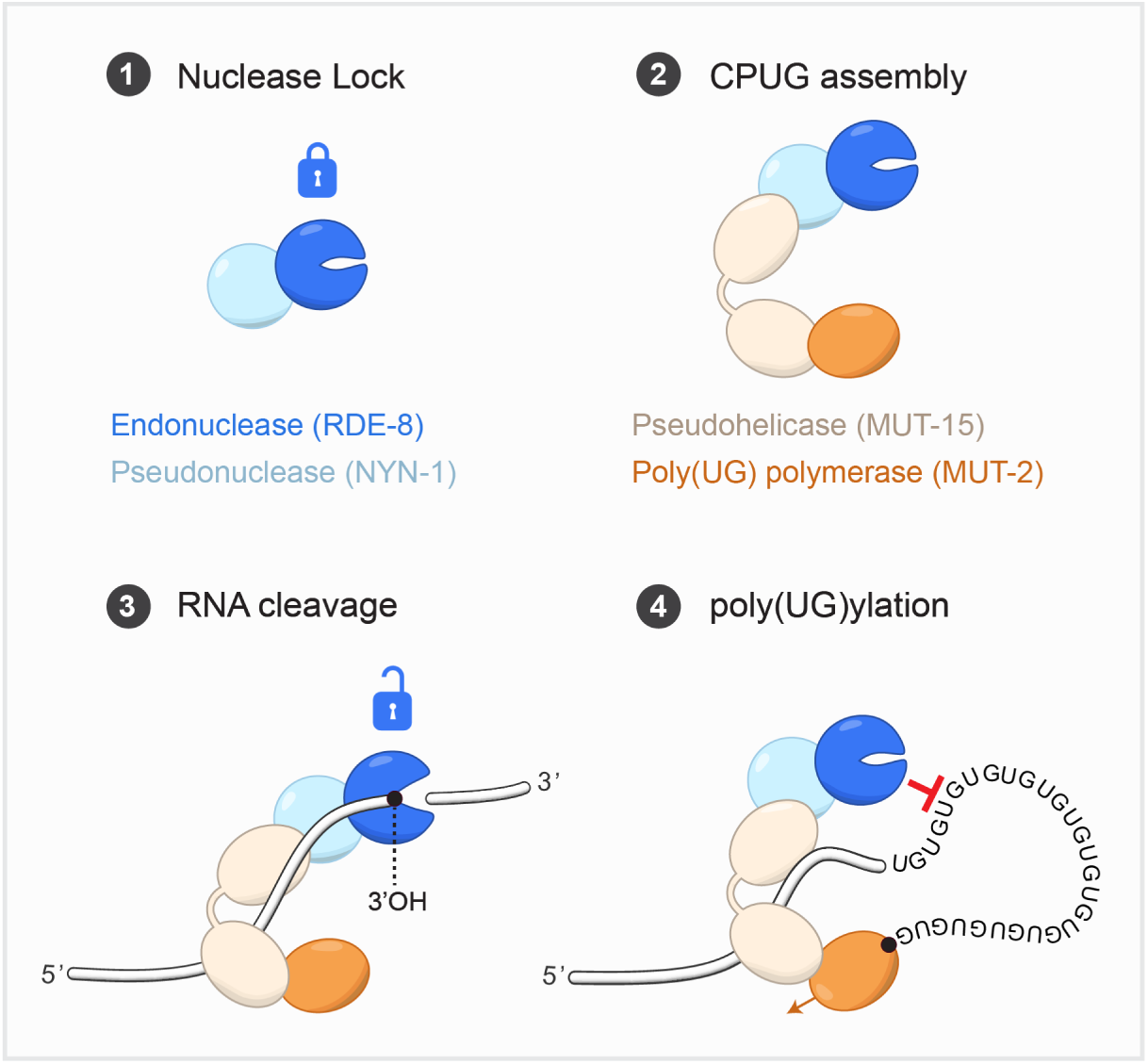

**Highlights:**

- RDE-8 forms an autoinhibited heterodimer with the pseudonuclease NYN-1
- MUT-15 is a pseudohelicase that licenses RDE-8 activity upon CPUG assembly
- CPUG synthesizes long poly(UG) tails influenced by the substrate 3′ end
- RDE-8 sequence specificity protects poly(UG) tails from cleavage

## Introduction

Small RNA (sRNA) silencing pathways are central regulators of gene expression, controlling transcript stability, translation, and chromatin states. These pathways are initiated by Argonaute proteins loaded with small RNAs, which recognize complementary target transcripts^1–3^. In *Caenorhabditis elegans*, primary target recognition is amplified through the production of secondary 22G sRNAs by RNA-dependent RNA polymerases (RdRPs)^4–6^. This sRNA amplification pathway operates downstream of multiple primary sRNA pathways, including exogenous RNAi, piRNAs and endogenous siRNAs, allowing diverse silencing signals to converge on a common amplification machinery. The resulting 22G RNAs are loaded into secondary Argonaute proteins^7^ that reinforce and maintain silencing of diverse targets^4,8^. Conversion of targeted transcripts into templates for sRNA amplification proceeds through two consecutive RNA-processing reactions. Following recognition by a primary Argonaute, the NYN-domain-containing endoribonuclease RDE-8, a member of the Regnase family^9,10^, cleaves target RNAs. The terminal ribonucleotidyltransferase MUT-2 (also known as RDE-3) then adds a poly(UG) tail to the newly generated 3′ end of the 5′ cleavage fragment^11–15^. These poly(UG) tails adopt G4 quadruplex-like structures that are resistant to the exonuclease MUT-7 and mark the RNA as a template for RdRP-dependent 22G sRNA synthesis (**Figure 1A**)^12,13,16–18^.

**Figure 1.**
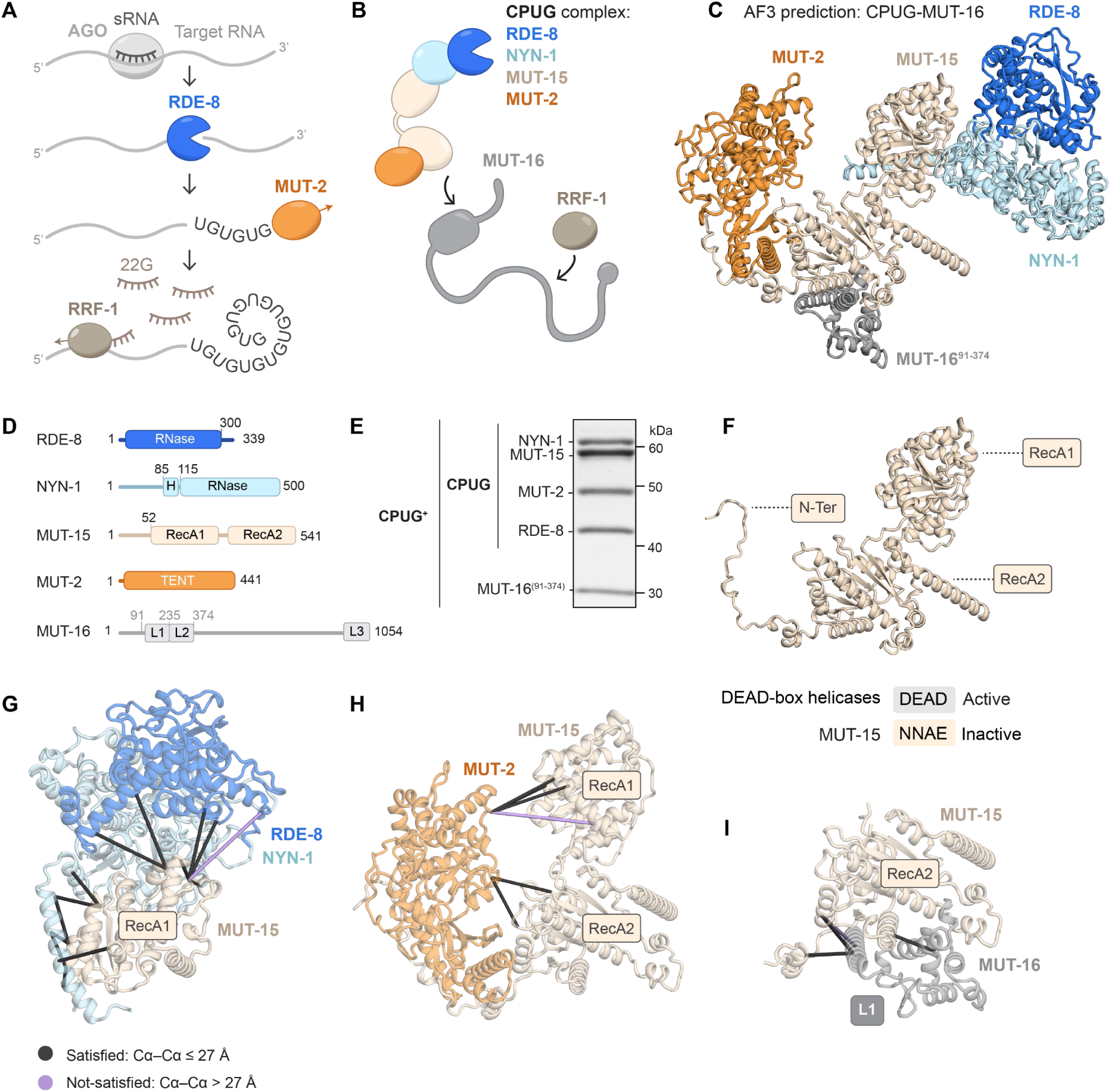
The CPUG complex is organized around a pseudohelicase. **(A**) Current model of small RNA amplification in *C. elegans*. **(B)** Schematic of the CPUG complex and its connection to the RNA-dependent RNA polymerase RRF-1 through MUT-16. **(C)** AlphaFold3 model of the CPUG complex bound to MUT-16, as described in Lowe et al.^16^. **(D)** Domain organization of the CPUG components and MUT-16. Construct boundaries used in this study are indicated. RNase=ribonuclease; H=α-helix; TENT=terminal nucleotidyltransferase; L1–L3=LOTUS 1–3. **(E)** SDS-PAGE and Coomassie staining of the purified CPUG^+^ complex. **(F)** AlphaFold3 model of MUT-15 with the RecA1- and RecA2-like domains indicated. Substitutions in the DEAD motif of MUT-15 compared with active DEAD-box helicases are highlighted. **(G–I)** Crosslinking analysis of the CPUG^+^ complex. Intermolecular crosslinks between MUT-15 and the RDE-8/NYN-1 nuclease module (G), MUT-2 (H), and the MUT-16 L1 domain (I) were mapped onto the AlphaFold3 model of CPUG^+^. See also Figure S1.

Cleavage and poly(UG)ylation are likely to require physical coupling to efficiently channel targeted RNAs into small RNA amplification. Using AlphaFold modeling together with *in vivo* co-immunoprecipitation experiments, the Kennedy group recently proposed that RDE-8 and MUT-2 are integrated into a four-protein complex, termed the pUGasome, together with the Regnase-family pseudonuclease NYN-1 (or its paralog NYN-2) and the protein of unknown domain structure MUT-15^16^. In the AlphaFold model, NYN-1 connects RDE-8 to MUT-15, whereas MUT-15 links the RDE-8/NYN-1 nuclease module to MUT-2. The study further proposed that the scaffold protein MUT-16 connects this machinery to the RdRP RRF-1, thereby linking RNA cleavage and poly(UG)ylation to downstream 22G-RNA synthesis (**Figure 1B,C**)^16^. While this study focused on the pUGasome role in antiviral RNAi in somatic cells, its components had previously been characterized primarily in the germline, where they localize to perinuclear Mutator foci and contribute to transposon silencing and thereby to genome integrity^9,19–22^.

These studies provided a plausible molecular architecture for physically coupling RNA cleavage and poly(UG)ylation. However, the pUGasome had not been biochemically reconstituted, and most of the predicted interaction surfaces remained experimentally untested^16^. Above all, it remained unknown how the activities of RDE-8 and MUT-2 are regulated and coordinated. Specifically, the complex must solve two key regulatory problems.

First, RDE-8 nuclease activity must be tightly controlled to prevent off-target RNA cleavage. Second, because RDE-8 cleaves RNA whereas MUT-2 extends it, these opposing activities must be coordinated to impose directionality, so that the poly(UG)-tailed template escapes re-cleavage and remains a substrate for RdRP-dependent amplification. Whether and how NYN-1 and MUT-15 contribute to these regulatory steps, beyond organizing the complex, remains unknown.

Here, we establish the first biochemical reconstitution of the pUGasome, which we refer to as the cleavage and poly(UG)ylation (CPUG) complex, and define the molecular principles underlying its regulation. The crystal structure of the isolated RDE-8/NYN-1 nuclease module reveals that the RDE-8 active site is locked in an autoinhibited conformation by a defined network of interactions. Biochemical analyses show that the nuclease module is inactive. We identify MUT-15 as a catalytically inactive DEAD-box-like pseudohelicase and show that its binding remodels the RDE-8/NYN-1 nuclease module, licensing RDE-8 activity upon CPUG complex assembly. In addition to supporting RNA cleavage, reconstituted CPUG synthesizes long, strictly alternating poly(UG) tails, and high-throughput sequencing defines how the substrate 3′ end specifies tailing efficiency and register. Finally, we identify two mechanisms that can impose directionality on the reaction: engagement of the RNA by MUT-2 limits its accessibility to RDE-8, while the cleavage preference of RDE-8 renders poly(UG) sequences intrinsically resistant to cleavage. Together, our findings show that RDE-8 activity is controlled by CPUG assembly and suggest how the interplay between RDE-8 and MUT-2 can impose directionality, safeguarding the RNA intermediate and directing it toward small-RNA amplification.

## Results

### The CPUG complex is organized around a pseudohelicase

The architecture of the cleavage and poly(UG)ylation (CPUG) complex, as well as its interaction with the scaffold protein MUT-16 that links it to the RdRP, were recently proposed based on AlphaFold modeling and *in vivo* interaction studies (**Figure 1C**)^16,19,20^. To validate the proposed molecular organization of this machinery and to investigate its biochemical activity and regulation, we set out to reconstitute the complex.

We co-expressed the four CPUG components (RDE-8, NYN-1, MUT-15 and MUT-2) together with the interacting structured region of MUT-16 (residues 91-374) in insect cells (**Figure 1D**; **Table S1**). Multistep purification yielded a homogeneous five-subunit complex, as confirmed by SDS-PAGE and mass spectrometry. Because the reconstituted CPUG complex also contains MUT-16^91–374^, we refer to it as CPUG^+^ throughout this study (**Figure 1E** and **S1A**).

Crosslinking mass spectrometry was consistent with the predicted architecture and positioned MUT-15 as the central interaction hub connecting the RDE-8/NYN-1 nuclease module, MUT-2 and MUT-16 (**Figure S1B**). Because MUT-15 is an uncharacterized factor with no previously annotated structural domains, we investigated its architecture to understand the structural basis of this central scaffolding function. Structure-based searches with Foldseek^23^ revealed that MUT-15 adopts the tandem RecA1–RecA2 architecture characteristic of ATP-dependent DEAD-box helicases^24,25^. Despite this structural similarity, the Q motif of MUT-15, which in active helicases recognizes the adenine base of ATP, is absent, and its other ATP-binding and hydrolysis motifs are degenerate, including the canonical DEAD motif, which is replaced by NNAE (**Figure 1F** and **S1C,D**). MUT-15 therefore represents a catalytically inactive DEAD-box-like pseudohelicase whose architecture has been repurposed to scaffold the cleavage and poly(UG)ylation machinery.

The AlphaFold3 model^16^ predicts that the MUT-15 RecA1-like domain binds NYN-1, forming the interface between MUT-15 and the RDE-8/NYN-1 nuclease module, which is supported by crosslinking mass spectrometry (**Figure 1G**). For MUT-2, the model predicts contacts primarily with the RecA2-like domain and the N-terminal region of MUT-15, while the RecA1-like domain is positioned nearby without direct contacts. Crosslinks mapped to both the RecA1- and RecA2-like domains (**Figure 1H**), in broad agreement with this prediction. Finally, multiple crosslinks were detected between the MUT-15 RecA2-like domain and MUT-16^91–374^ (**Figure S1E**).

Structural analysis of this region identified two consecutive extended LOTUS (eLOTUS) domains (**Figure S1F**). AlphaFold3 predictions and crosslinking indicate that eLOTUS1 (residues 116-235) binds the RecA2-like domain of MUT-15 (**Figure 1I** and **S1G**), closely resembling the interaction between the eLOTUS domain of Oskar and the RecA2 domain of the DEAD-box ATPase Vasa from *Drosophila melanogaster*^26,27^ (**Figure S1H**). In contrast, eLOTUS2 (residues 239-374) is not predicted to contact MUT-15 directly but crosslinks with several CPUG components, consistent with a more flexible orientation (**Figure S1E,G**). Full-length MUT-16 contains a third, C-terminal minimal LOTUS (mLOTUS) domain that lacks the α-helix associated with DEAD-box protein binding and is not predicted to contact MUT-15 (**Figure S1F**).

Together, these data identify MUT-15 as a catalytically inactive pseudohelicase and provide a structural basis for its scaffolding role within CPUG. Despite the loss of catalytic activity, MUT-15 retains the RecA2 surface that active DEAD-box helicases use to bind eLOTUS domains, and engages MUT-16 through this conserved interface, thereby connecting CPUG to the Mutator scaffold.

### An active-inactive Regnase pair forms the nuclease module of CPUG

The catalytically active nuclease RDE-8 associates with the pseudonuclease NYN-1 to form a stable heterodimer^16^. RDE-8 and NYN-1 both belong to the Regnase family of NYN-domain ribonucleases, which have important roles in cellular defense across animals^10,28,29^. In mammals, Regnase-1 forms homodimers, and dimerization has been implicated in its RNase activity^28,30,31^. The unusual pairing of an active nuclease with an inactive family member therefore raised the question of how heterodimerization affects RDE-8 structure and catalytic state. We addressed this by determining the structure of the RDE-8/NYN-1 nuclease module.

To this end, we generated minimal RDE-8 and NYN-1 constructs that encompass the NYN domains^32^ and the folded flanking regions of both proteins. We determined the crystal structure of RDE-8/NYN-1 in the presence and absence of Mg^2+^ or Mn^2+^ ions. All structures were highly similar, indicating that the overall architecture and active-site conformation are insensitive to the presence of Mg^2+^ or Mn^2+^. We therefore report the highest-resolution structure (1.65 Å) as the representative model (**Figure 2A**; **Table S2**). The nuclease RDE-8 binds NYN-1 through an extended interface spanning ∼1700 Å^2^. The RDE-8/NYN-1 heterodimer adopts a NYN-domain arrangement similar to that observed in crystal structures of mammalian Regnase family homodimers^30,33–35^, with additional flanking regions that pack against the NYN core and extend its architecture (**Figure S2A,B**).

**Figure 2.**
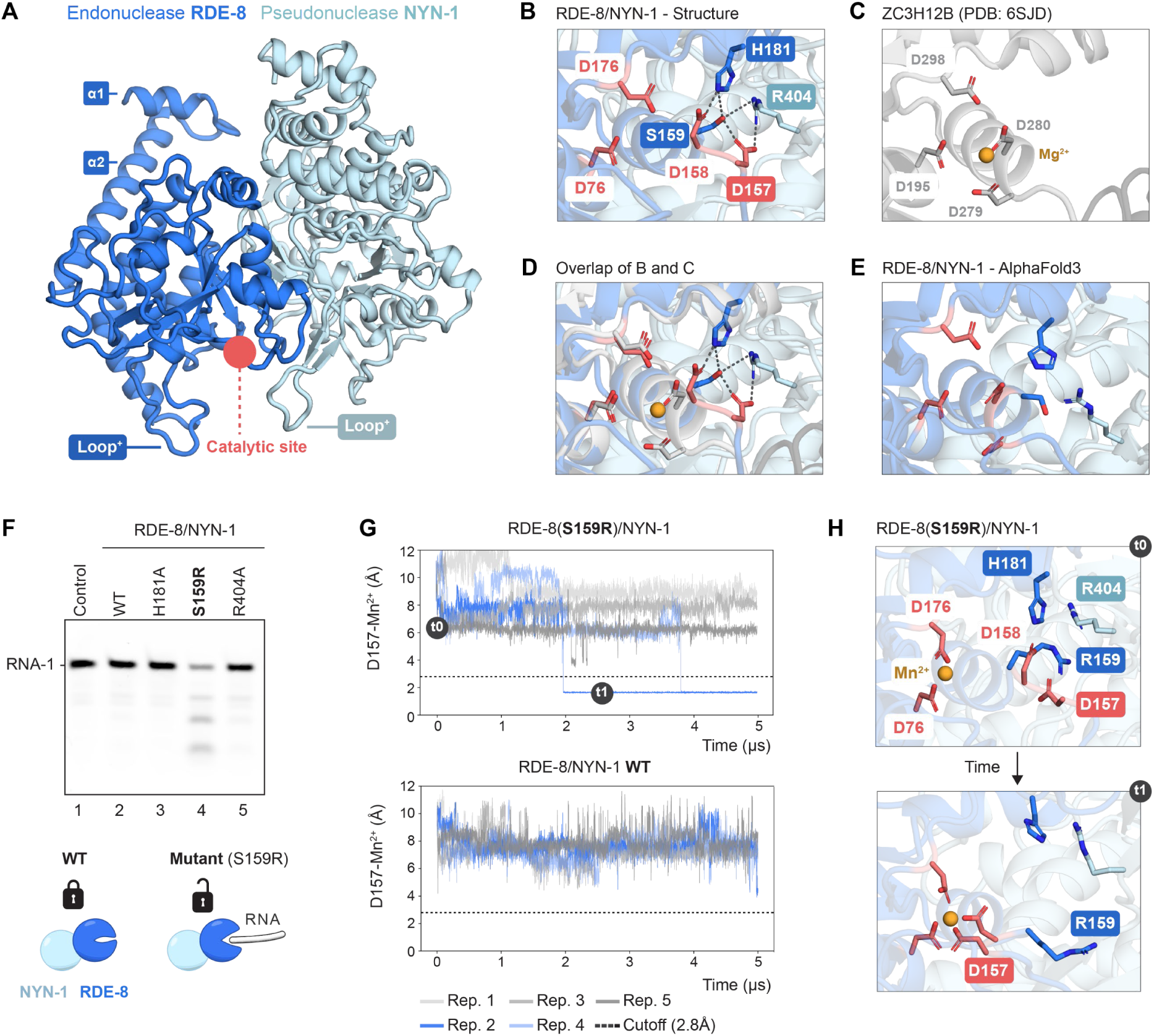
An inhibitory network restrains RDE-8 nuclease activity (A-B) Crystal structure of the RDE-8/NYN-1 complex determined in this study (PDB: 28RS). Positive loops Loop^+^) are indicated. **(B)** Zoom-in of RDE-8 active site from the structure shown in (A), highlighting the catalytic aspartates and residues forming the inhibitory network. Dashed lines indicate interactions between residues. **(C)** Zoom-in of the active site of human ZC3H12B (PDB: 6SJD)^35^. **(D)** Superposition of (B) and (C). **(E)** Zoom-in of RDE-8 active site in the RDE-8/NYN-1 AlphaFold3 model. **(F)** Nuclease assays using wild-type (WT) or mutant RDE-8/NYN-1 complexes and a 5′-FAM-labeled 16-nt RNA substrate (RNA-1). Control: RNA without protein. Representative of three independent experiments. **(G,H)** Molecular dynamics (MD) simulations of WT or mutant (S159R) RDE-8/NYN-1. The distance between RDE-8 D157 and the Mn²⁺ ion is plotted over time (G). Structural snapshots from one RDE-8(S159R)/NYN-1 simulation illustrating the rearrangement of the catalytic aspartates into the active conformation (H). See also Figure S2, Table S2 and Table S3.

To understand how the RDE-8/NYN-1 complex differs from mammalian Regnase-family members, we generated a structure-based sequence alignment of human and *C. elegans* Regnase-family proteins using the AlphaFold3 models of their NYN domains (**Figure S2C**). Although the overall fold is conserved, several loop regions have diverged substantially. The alignment revealed two main features that distinguish the RDE-8/NYN-1 heterodimer. First, the canonical RNA-binding loop of mammalian Regnases is absent from both RDE-8 and NYN-1 (**Figure S2D**). Instead, the heterodimer carries two positively charged insertions that flank the catalytic cleft. A NYN-1-specific loop occupies approximately the position of the mammalian RNA-binding loop, whereas an RDE-8-specific insertion forms the opposite wall of the cleft (**Figure 2A** and **S2A**). Second, mammalian Regnases homodimerize through a conserved dimerization loop^30^. NYN-1 retains a recognizable dimerization loop, whereas RDE-8 carries a large insertion at this position (**Figure S2C**). This difference may help explain why RDE-8 and NYN-1 associate as a heterodimer rather than forming homodimers, as seen for the mammalian Regnases.

To understand how partner specificity is achieved within the *C. elegans* Regnase family, we used AlphaFold3^36^ to predict pairwise interactions among its members. Of the proteins analyzed, only RDE-8 and NYN-3 retain the catalytic residues characteristic of active nucleases, whereas the others have degenerate active sites^10^ (**Figure S2E**). Across the predictions, both active nucleases were predicted to pair with specific inactive family members rather than to homodimerize. RDE-8 was predicted to dimerize with NYN-1, NYN-2 and ERI-9, whereas NYN-3 was predicted to interact with the uncharacterized paralogs Y24D9B.1 and Y51H4A.13 (**Figure S2F**). Notably, RDE-8 and ERI-9 have both been linked to ERIC-mediated 26G-sRNA biogenesis during oogenesis and embryogenesis^9,37,38^, whereas NYN-3 functions in ERIC-mediated 26G-sRNA biogenesis during spermatogenesis^37,39,40^. Comparison of the predicted complexes revealed a shared heterodimerization mode (**Figure S2G**). Interaction with MUT-15 was predicted specifically for the RDE-8/NYN-1 pair, with NYN-1 mediating the contact (**Figure S2H**). Together, these predictions suggest that pairing an active with an inactive Regnase is a recurring feature across distinct small-RNA pathways, and that inactive Regnase-family partners may contribute to pathway specificity by directing active nucleases into distinct protein assemblies.

### The isolated nuclease module adopts an autoinhibited conformation

Inspection of the catalytic center of RDE-8 in our crystal structure unexpectedly revealed an inactive conformation (**Figure 2B**). Comparison with the human Regnase-1 family member ZC3H12B^35^ showed that two of the four residues that form the active site of RDE-8, D157 and D158, are rotated away from the productive arrangement required for metal ion coordination and catalysis (**Figure 2C,D**). Notably, AlphaFold3 predicts the catalytic aspartates of RDE-8 to adopt the same active conformation observed for ZC3H12B and other mammalian members of the family (**Figure 2E**), indicating that the autoinhibited state identified experimentally is not captured by current structure prediction methods. The inactive orientation is stabilized by a network of intra- and inter-molecular interactions involving RDE-8 S159 and H181 together with NYN-1 R404, which belongs to the NYN-1/NYN-2-specific positive loop (**Figure S2C**). H181 forms a hydrogen bond with D158, while D157 is contacted by both S159 and R404. S159 occupies a central position in this network, contacting D157, H181, and R404 simultaneously, and R404 further staples these residues together by contacting both D157 and H181 (**Figure 2B**). This conformation is unlikely to result from the absence of metal ions, since Mg^2+^ and Mn^2+^ were present during crystallization, but none were observed in the active site. Co-crystallization or soaking with short RNAs under multiple conditions yielded no RNA density and did not alter this state, leaving all structures indistinguishable from the apo complex. RDE-8/NYN-1 therefore appears to adopt an intrinsically autoinhibited conformation. Consistent with a functional role, residues S159, H181 and R404 are highly conserved throughout *Caenorhabditis* (**Figure S2I**).

### Disruption of the autoinhibitory network enables RNA cleavage

To determine whether the autoinhibited conformation restrains RDE-8 nuclease activity, we purified full-length RDE-8/NYN-1 complexes, either wild-type or carrying mutations within the inhibitory network (RDE-8 S159R, RDE-8 H181A, and NYN-1 R404A) and compared their endonuclease activities. H181A and R404A are alanine substitutions that remove the side chains contributing to the inhibitory network. In contrast, S159R replaces the serine found in *C. elegans* RDE-8 with the arginine present at this position in mammalian Regnases (**Figure S2C**). A fluorescently labeled 16-nt RNA (RNA-1) was used as substrate. Neither the wild-type RDE-8/NYN-1 complex nor RDE-8 alone generated detectable cleavage products. RDE-8/NYN-1 purified from insect cells was also inactive, indicating that the lack of activity is not a consequence of bacterial expression (**Figure 2F** and **S2J**). Whereas the RDE-8(H181A)/NYN-1 and RDE-8/NYN-1(R404A) mutant complexes remained inactive, RDE-8(S159R)/NYN-1 displayed robust activity (**Figure 2F**). RDE-8 activity was strictly dependent on Mn^2+^, and no cleavage was observed in the absence of divalent cations or in the presence of Mg^2+^ (data not shown). Thus, disrupting the center of the inhibitory network by introducing arginine in place of S159 is sufficient to activate RDE-8, whereas removing individual peripheral contacts is not.

To determine whether the activating S159R mutation stabilizes the active conformation of RDE-8, we determined the crystal structure of the RDE-8(S159R)/NYN-1 complex in the presence of Mn^2+^ at 2.0 Å resolution (**Table S2**). Surprisingly, despite its robust nuclease activity, the catalytic aspartates again adopted the inactive conformation observed in the wild-type structure, and no Mn^2+^ ion was bound at the active site despite its presence during crystallization (**Figure S2K**). Nevertheless, the structure revealed two localized conformational changes in the inhibitory network. The RDE-8 loop directly following H181 (residues 183-187) rearranges, and NYN-1 R404 undergoes a rotamer change together with a shift of the adjacent loop. Together, these changes break the trans-contact that R404 makes to RDE-8 in the wild type and partially open the catalytic cleft. Thus, although the catalytic aspartates remained displaced, S159R disrupts the inter-subunit network that stabilizes autoinhibition, which may favor a shift toward the active state.

To test this possibility, we performed molecular dynamics (MD) simulations of the RDE-8/NYN-1 wild-type and mutant (S159R) complexes, running five independent replicates of 5 μs each with the Mn^2+^ ion placed near the catalytic aspartates. In all wild-type simulations, D157 always remained ≥3.5 Å from the metal throughout the simulations. In two of five S159R simulations, however, D157 rotated from its tilted, autoinhibited conformation into the canonical active orientation and directly coordinated the Mn^2+^ ion. (**Figure 2G**). Once formed, the D157-Mn^2+^ coordination persisted for the remainder of the trajectory in both cases, with no return to the tilted state (**Figure 2G, S2L; Table S3**). The active conformation is therefore reached only rarely but is stable once attained, consistent with the S159R substitution permitting access to a catalytically competent state that is itself intrinsically stable. D158 coordinated the ion stably in all simulations of both wild-type and mutant, indicating that the S159R substitution acts specifically on D157 (**Figure S2L**). Adoption of the active conformation correlated with displacement of the NYN-1 positive loop carrying R404. In this displaced conformation, R404 is no longer engaged in the inhibitory network, and this region of the cleft is exposed, potentially making it available for RNA binding. Notably, in the S159R crystal structure, this loop also shifts position, indicating that the structure might capture an early stage of activation rather than the fully active state (**Figure S2M**).

Together, these findings indicate that the inactive conformation identified in the crystal structure of the wild-type complex is actively maintained by a network of intra- and intermolecular interactions that restrain RDE-8 activity. Disrupting this network increases access to, but does not fully stabilize, the active conformation, implying that additional factors might be required for full catalytic activation.

### Activation by MUT-15 uncovers RDE-8 cleavage preference

Having established that RDE-8/NYN-1 is intrinsically autoinhibited, but knowing its activity is necessary for 22G sRNA biogenesis^9^, we then asked whether an additional factor is required to activate the nuclease. The pseudohelicase MUT-15 was a strong candidate, since it binds RDE-8/NYN-1 and serves as the structural scaffold of the CPUG complex.

Attempts to purify MUT-15 alone were unsuccessful. We therefore removed its disordered, hydrophobic N-terminal region (residues 1-51), which binds MUT-2, and co-expressed MUT-15^52–541^ with MUT-16^91–374^. This yielded a stable MUT-15^52–541^/MUT-16^91–374^ subcomplex. Adding MUT-15^52–541^/MUT-16^91–374^ to RDE-8/NYN-1 triggered RDE-8 nuclease activity. Comparable activity was seen for the co-purified RDE-8/NYN-1/MUT-15^52–541/MUT–1691–374^ complex, whereas complexes containing catalytically inactive RDE-8 mutants (D76N or D158N) produced no detectable cleavage products (**Figure 3A** and **S3A**). Nuclease activity was strictly dependent on Mn^2+^, and no cleavage was observed in the presence of Mg^2+^ (**Figure 3B**). Together, these data show that RDE-8/NYN-1 nuclease activity is unlocked by MUT-15. We hereafter refer to the minimal active four-protein assembly (RDE-8/NYN-1/MUT-15^52–541^/MUT-16^91–374^) as the RDE-8 cleavage complex (RDE-8^CC^).

**Figure 3.**
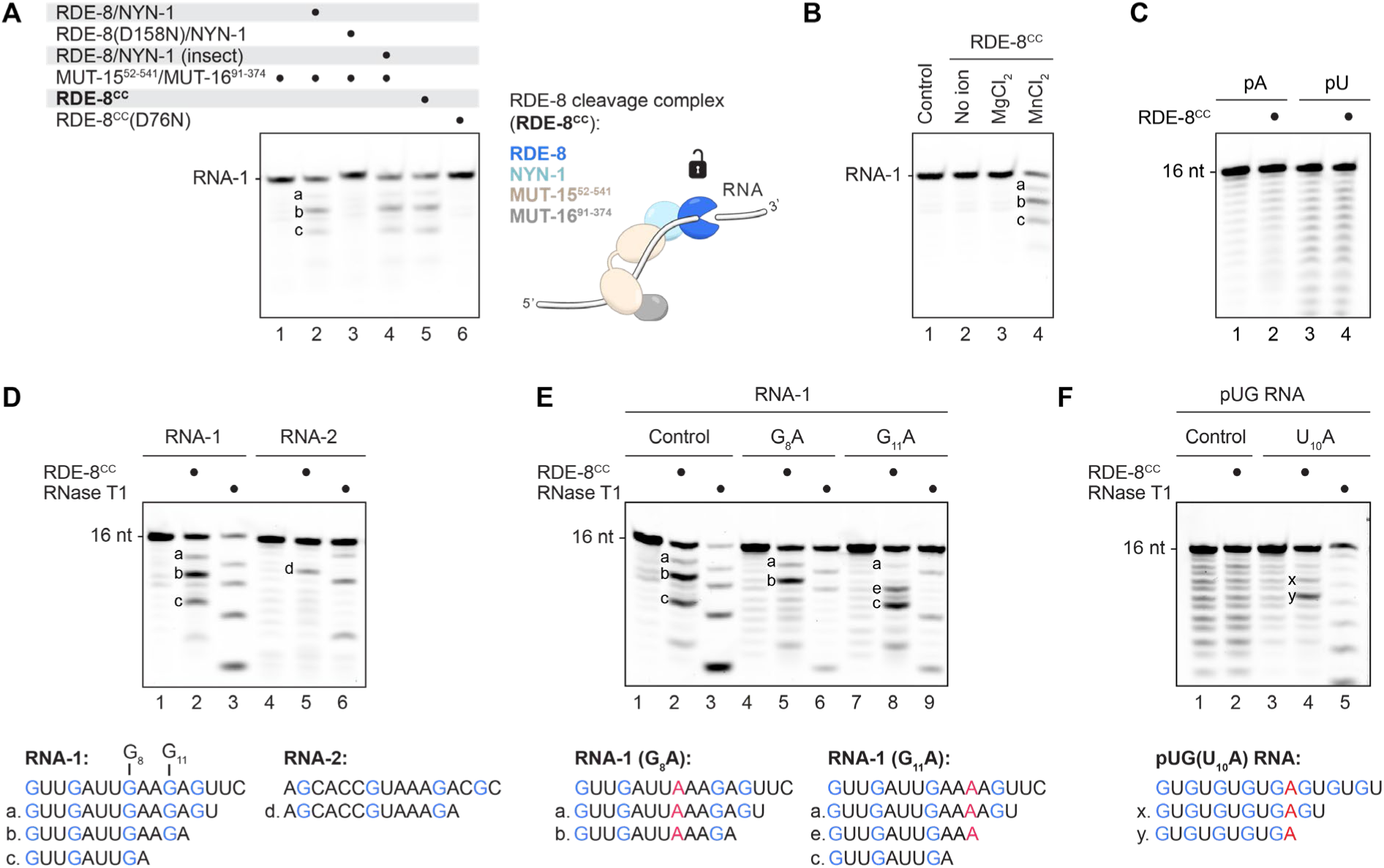
Activation by MUT-15 uncovers RDE-8 cleavage preference (A–F) RDE-8-mediated RNA cleavage was assessed using the indicated protein complexes, and 16-nt 5′-FAM-labeled RNA substrates. Representative of three independent experiments. **(A)** Wild-type (WT) or mutant RDE-8/NYN-1 mixed with MUT-15^52–541/MUT–1691–374^, and the co-purified RDE-8^CC^ tested with substrate RNA-1. Catalytically inactive complexes (RDE-8 D76N or D158N) served as negative controls. **(B)** Metal dependence of RDE-8^CC^. Cleavage assays were performed with RNA-1 in the presence of Mn²⁺, Mg²⁺, or no divalent cation. **(C)** Substrate preference of RDE-8^CC^ assessed using poly(A) (pA) and poly(U) (pU) RNA substrates. **(D)** Identification of RDE-8^CC^ cleavage products generated from the RNA-1 and RNA-2 substrates. RNase T1, which cleaves 3′ to guanosine residues, was included as a reference. Cleavage products are labeled a–c for RNA-1 and d for RNA-2. **(E)** Effect of the G_8_A or G_11_A substitutions on the RNA-1 (Control) cleavage pattern. Cleavage products are labeled as in (D). **(F)** Cleavage of a 16-nt pUG RNA and a version carrying a U_10_A substitution. Cleavage products are labeled as x and y. See also Figure S3.

Cleavage of the 16-nt RNA substrate (RNA-1) by RDE-8^CC^ yielded three discrete products rather than the smear expected from complete or nonspecific degradation (**Figure 3A**,**B**). Moreover, 16-nt poly(U) or poly(A) RNAs were not cleaved, suggesting that RDE-8 cleaves RNA with a preference for certain nucleotide compositions (**Figure 3C** and **S3B**). We thus sought to define the cleavage specificity of RDE-8^CC^. To map the cleavage sites for RNA-1, we used RNase T1, which specifically cleaves 3′ to guanosine nucleotides, as a reference. The three RDE-8^CC^ products migrated one nucleotide slower than the corresponding RNase T1 bands. A product one nucleotide longer therefore indicates that RDE-8^CC^ cleaves one position further along, leaving a GN dinucleotide at the 3′ end of the 5′ cleavage fragment. A second 16-nt RNA substrate (RNA-2) similarly yielded a cleavage product terminating in GA (**Figure 3D** and **S3B**). Replacement of the guanosine at position 8 (G_8_) with adenosine in RNA-1 abolished generation of the fragment terminating at G_8_A_9_ (fragment c), supporting the importance of this guanosine. Replacing G_11_ with adenosine similarly resulted in the loss of the fragment terminating at G_11_A_12_ (fragment b) but instead produced a new cleavage product terminating at A_10_A_11_ (fragment e) (**Figure 3E** and **S3C**).

These results indicate that guanosine contributes to RDE-8 cleavage-site selection. If guanosine alone specified cleavage, an alternating GU sequence would provide numerous potential cleavage sites. This is particularly relevant because MUT-2 synthesizes poly(UG) tails, which must remain intact to support downstream sRNA amplification. We therefore asked whether GU-repeat RNA is susceptible to RDE-8 cleavage. To test this, we used a 16-nt RNA consisting of eight GU repeats (pUG RNA), which is too short to adopt the characteristic pUG fold^13^. While we observed no cleavage for this pUG RNA, a single substitution of a uridine to adenosine (U_10_A) resulted in cleavage. Cleavage again occurred one nucleotide downstream of guanosine, generating a main cleavage product terminating in UGA and a weaker one in AGU (**Figure 3F** and **S3C**).

Together, these findings indicate that although guanosine contributes to RDE-8 cleavage-site selection, cleavage also depends on the surrounding sequence context, with repetitive UG sequences being intrinsically resistant to cleavage. This suggests that poly(UG) tails synthesized by MUT-2 are protected from RDE-8-mediated re-cleavage during 22G-RNA biogenesis.

### MUT-15 remodels the nuclease module upon CPUG assembly

To understand how MUT-15 activates RDE-8, we sought to determine the structure of RDE-8/NYN-1 bound to MUT-15. Crystallization trials did not yield crystals, so we turned to cryo-EM and determined the structure of the reconstituted CPUG^+^ complex. In single-particle cryo-EM analysis, we observed 2D classes that were in agreement with the intact complex, but several classes indicated strong flexibility (**Figure S4A**). This conformational heterogeneity is consistent with the AlphaFold3 predictions (**Figure S4B**) and crosslinking mass spectrometry data (**Figure S1E**). Despite attempts to overcome this issue using different refinement strategies, we were only able to obtain high-resolution density for a rigid subcomplex, resulting in a 2.9 Å map comprising the RDE-8/NYN-1 module bound to the RecA1-like domain of MUT-15 (**Figure 4A** and **S4C-J**; **Table S4**). The remaining elements, the RecA2-like domain of MUT-15, MUT-2 and MUT-16, were not resolved and showed only weak density, indicating that they are flexibly tethered to this ordered core.

**Figure 4.**
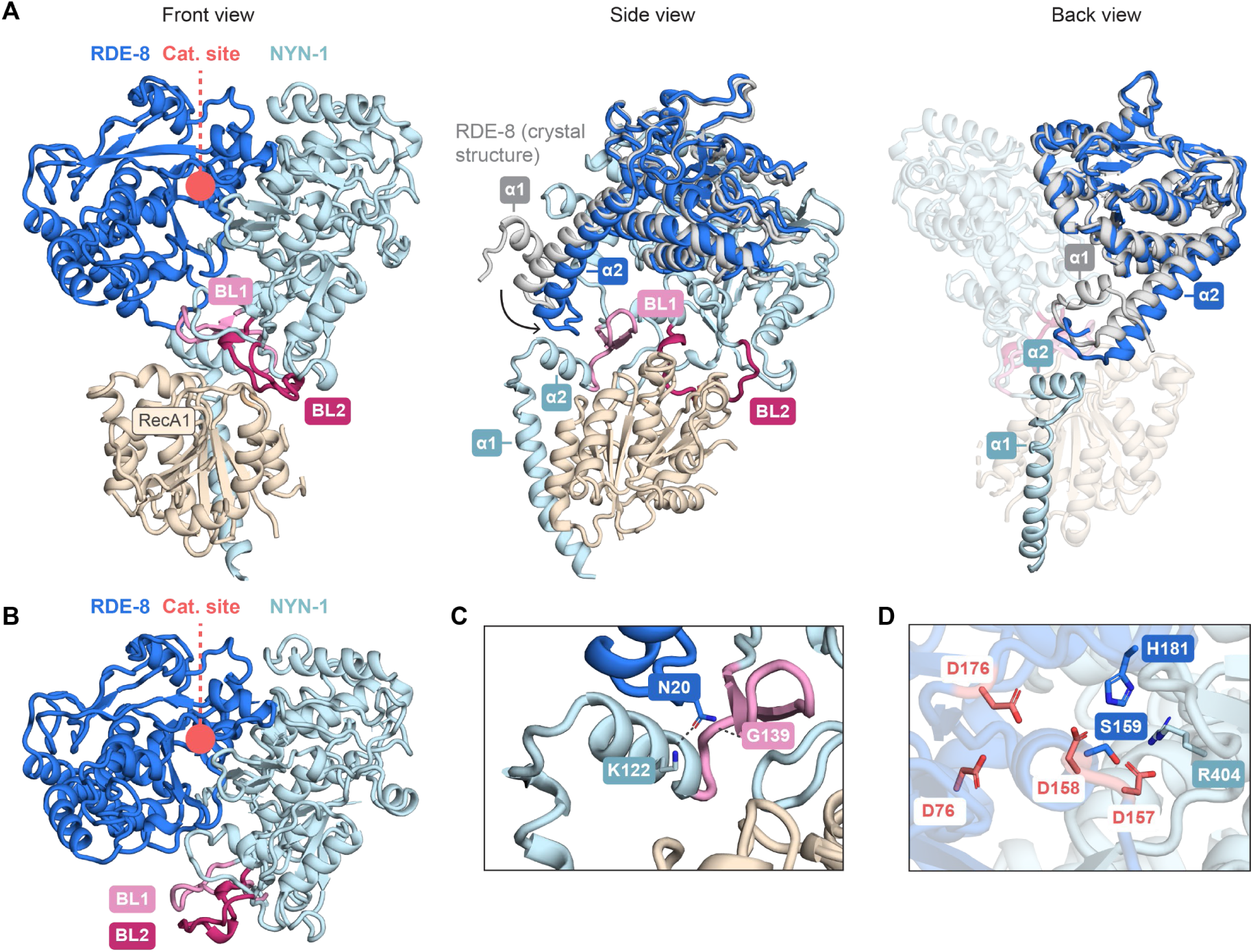
MUT-15 remodels the nuclease module without relieving autoinhibition. **(A**) Cryo-EM structure of RDE-8/NYN-1/MUT-15^RecA1^ shown in three orientations (PDB: 32VT). In the side and back views, RDE-8 from the RDE-8/NYN-1 crystal structure (aligned on NYN-1) is shown in grey to illustrate the conformational change associated with MUT-15 binding. RDE-8^α1^ could be modeled only in the crystal structure because of the low local quality of the cryo-EM map. **(B)** Crystal structure of the RDE-8/NYN-1 complex determined in this study (PDB: 28RS). NYN-1^BL1^ and NYN-1^BL2^ are highlighted as in (A) to illustrate their conformational rearrangement upon MUT-15 binding. **(C)** Zoom-in view from (A) of the interaction of RDE-8 with NYN-1^α2^ and NYN-1^BL1^. **(D)** Zoom-in view of the RDE-8 catalytic site from (A). See also Figure S4, Table S4 and Movie S1.

The RecA1-like domain of MUT-15 docks onto NYN-1 across an extended interface of approximately 1,700 Å² formed by two contacts. The N-terminal region of NYN-1 folds into an L-shaped element formed by two alpha helices (NYN-1^α1^, NYN-1^α2^) connected by a short turn, with NYN-1^α1^ (residues 88-111) binding directly in a hydrophobic groove on the RecA1-like domain. An additional interaction interface consists of two aromatic-rich binding loops (NYN-1^BL1^, NYN-1^BL2^) present in the NYN-1 core (residues 128-143, 271-278) that pack directly against a complementary hydrophobic surface on MUT-15 RecA1 (**Figure 4A** and **S4K**). To test the contribution of NYN-1^α1^ to MUT-15 binding, we performed pull-down assays using different RDE-8/NYN-1 constructs. Deletion of the disordered NYN-1 N-terminal region (residues 1-84) did not affect association with MUT-15^52–541/MUT–1691–374^, whereas additional removal of NYN-1^α1^ (residues 1-114) abolished complex formation, demonstrating that this helix is required for binding of the nuclease module to the pseudohelicase scaffold (**Figure S4L**).

To determine how MUT-15 binding affects the nuclease module, we compared the cryo-EM structure with the RDE-8/NYN-1 crystal structure. Superposition of the two structures revealed two principal differences. First, in the crystal structure, NYN-1^BL1^ and NYN-1^BL2^ adopt a conformation that is incompatible with MUT-15 binding (**Figure 4B**). In the cryo-EM structure, both loops rearrange to accommodate the RecA1-like domain (**Figure 4A**, front view). Second, MUT-15 binding is accompanied by a rotation of RDE-8 relative to NYN-1. A region of RDE-8 N-terminal to its NYN domain forms a U-shaped element comprising RDE-8^α1^ (residues 7-13) and RDE-8^α2^ (residues 23-47), connected by a short linker (**Figure 4A**, back view). This U-shaped element rotates by approximately 11° upon MUT-15 binding, accompanied by an approximate 4° rotation of the RDE-8 NYN-domain core (**Figure 4A**, side view). The linker within the RDE-8 U-shaped element contacts NYN-1^α2^, with RDE-8 N20 positioned between NYN-1 K122 and G139 (**Figure 4C**). This interlocking contact connects RDE-8 to regions of NYN-1 that rearrange to accommodate MUT-15 (**Figure 4A**, side view), providing a potential pathway through which MUT-15 docking could promote reorientation of the RDE-8 NYN domain (Movie S1).

Despite these rearrangements, the RDE-8 active site retains the inactive conformation observed in the crystal structure (**Figure 4D**). Thus, although CPUG^+^ is active *in vitro*, the cryo-EM structure does not capture a catalytically competent nuclease state. This mirrors the crystal structure of the biochemically active RDE-8(S159R)/NYN-1 mutant, in which the active site likewise remained in an inactive conformation. MUT-15 binding might therefore increase access to a catalytically competent state without releasing the autoinhibited active site.

Completion of this transition likely requires binding of a metal ion and the RNA substrate. MUT-15 has an extensive positively charged patch on its surface^16^ (**Figure S4M**), suggesting that the pseudohelicase could bind RNA within the complex. Consistent with this, fluorescence polarization measurements showed that MUT-15^52–541^/MUT-16^91–374^ bound a 16-nt RNA substrate with high affinity (*K*_d_ = 0.26 μM), whereas MUT-16^91–374^ alone showed no detectable binding (**Figure S4N**). As expected from MUT-15’s degenerate ATP-binding and hydrolysis motifs, purified MUT-15^52–541^/MUT-16^91–374^ showed no detectable ATPase activity, supporting the conclusion that MUT-15 is a catalytically inactive pseudohelicase (**Figure S4O**).

Together, these results show that the pseudohelicase MUT-15 docks onto the RDE-8/NYN-1 dimer and remodels its surface while the autoinhibited state of the active site is retained. By providing an RNA-binding platform, MUT-15 may help recruit and position the RNA substrate within CPUG, ultimately leading to RDE-8 activation and RNA cleavage.

### RNA cleavage and tailing are coordinated within CPUG

The terminal nucleotidyltransferase MUT-2 synthesizes untemplated poly(UG) tails, but biochemical characterization of the *C. elegans* enzyme has been hindered because it cannot be purified in isolation (this study and ^14^). We therefore asked whether MUT-2 is catalytically active within the reconstituted CPUG^+^ complex.

To this end, we incubated purified CPUG^+^ with a fluorescently labeled 28-nt single-stranded RNA and individual ribonucleoside triphosphates (rNTPs), a UTP/GTP mixture, or all four rNTPs. Reactions were performed in the presence of Mg²⁺, which does not support RDE-8-mediated cleavage. When ATP or CTP were supplied, 3′-extensions of only one or two nucleotides were observed, indicating that MUT-2 initiates but does not elongate oligo(A) or oligo(C) tails. In contrast, reactions containing either GTP or UTP generated longer products, indicating more efficient elongation of G and U tails. Tailing reactions proceeded more efficiently in the presence of both UTP and GTP, or all four rNTPs, with products reaching ∼100 nt after 30 s and up to ∼300 nt after 5 min (**Figure 5A** and **S5A**).

**Figure 5.**
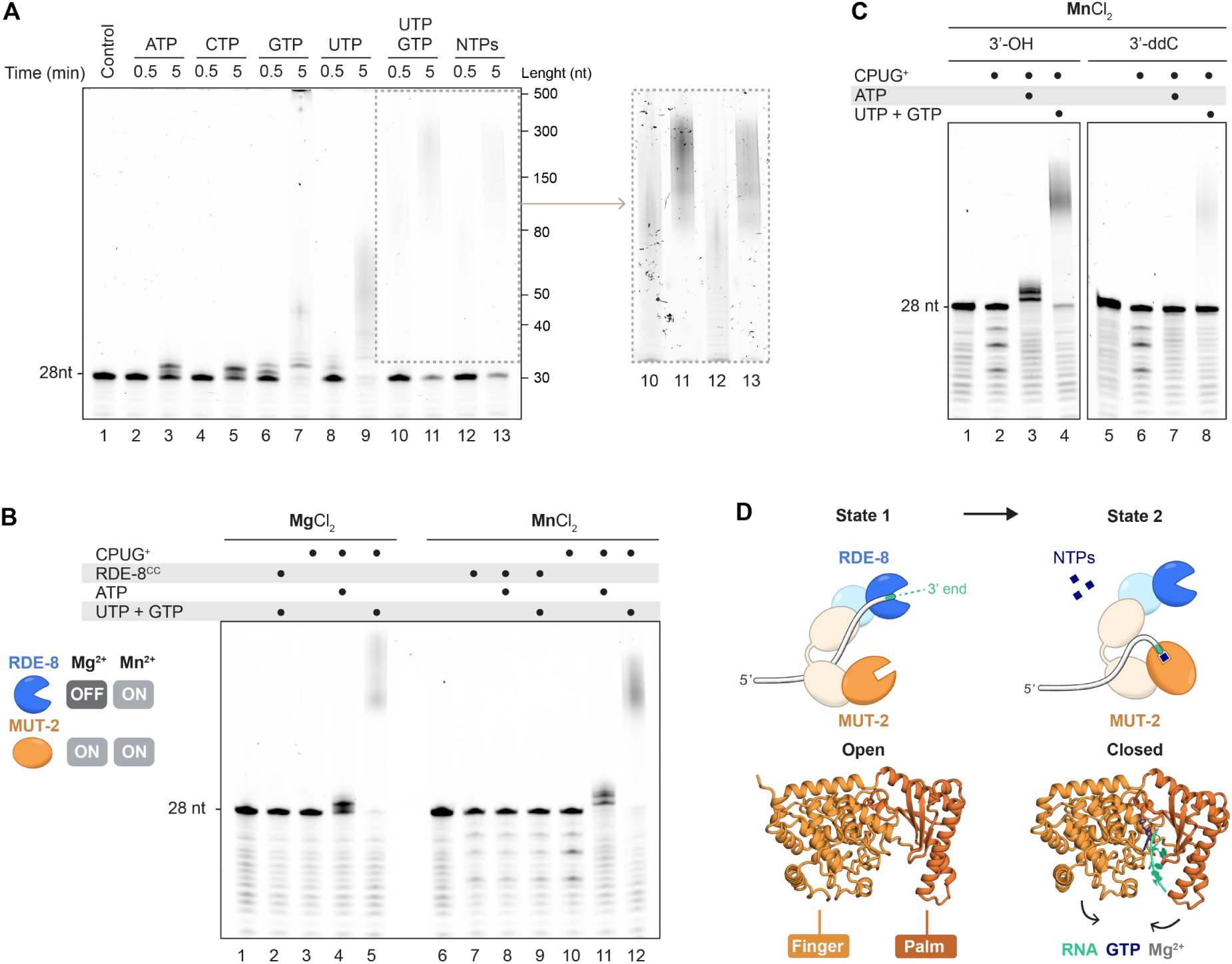
RNA cleavage and tailing are coordinated within CPUG (A–C) RNA cleavage and tailing assays using CPUG^+^ and 5′-FAM-labeled 28-nt RNA substrates. Representative of three independent experiments. **(A)** Nucleotide-incorporation preference of MUT-2. CPUG^+^ was incubated with individual rNTPs, a mixture of UTP and GTP, or all four rNTPs in Mg²⁺-containing buffer for 30 sec or 5 min. A higher-contrast view of the indicated region is shown. **(B)** Divalent-cation dependence of RDE-8-mediated cleavage and MUT-2-mediated tailing. CPUG^+^ was incubated in the presence of Mg²⁺ or Mn²⁺, with or without rNTPs. The RDE-8^CC^, which lacks MUT-2, was included as a control. **(C)** Comparison of an RNA carrying either a 3′-OH or a dideoxycytidine (3′-ddC). **(D)** Model of the mutually exclusive cleavage (state 1) and tailing (state 2) states of CPUG^+^, with MUT-2 modeled by AlphaFold3 in open (state 1) and closed (state 2) conformations^41^. Finger and palm domains are indicated. See also Figure S5.

To investigate the interplay between cleavage and tailing, we next performed reactions in the presence of Mn^2+^, which is permissive for RDE-8 activity. In the absence of rNTPs, CPUG^+^ produced RNA cleavage products characteristic of RDE-8 activity, similar to the RDE-8^CC^ complex, which was used as a control. As observed above, addition of ATP resulted in short tails, whereas UTP and GTP produced long tails, indicating that MUT-2 is also catalytically active with Mn^2+^. Notably, cleavage products were no longer detectable in the presence of rNTPs (**Figure 5B** and **S5B**). This observation could reflect either rapid tailing of cleavage products or preferential extension of the intact RNA before cleavage can occur.

To further investigate this, we analyzed an equivalent 28-nt RNA substrate terminating in a 3′-dideoxycytidine (3′-ddC), which lacks a 3′-hydroxyl group (3′-OH) and cannot be directly extended by MUT-2 without previous cleavage. In the absence of rNTPs, the 3′-ddC RNA was cleaved with efficiency comparable to that of the control RNA bearing a 3′-OH. Upon addition of ATP, the 3′-ddC RNA remained untailed, and, in the presence of UTP and GTP, little to no tailing was observed (**Figure 5C** and **S5C**). The strong reduction in tailing of the 3′-ddC substrate indicates that most tails observed for the 3′-OH RNA arise from direct extension of its pre-existing 3′ end rather than from extension of cleavage-generated products. Moreover, the loss of detectable cleavage products for both RNAs in the presence of rNTPs indicates that RDE-8-mediated cleavage is strongly suppressed under these conditions.

Together, these observations support a model in which CPUG^+^ exists in two states. In the absence of rNTPs, RNA can access the RDE-8 active site for cleavage. In contrast, when rNTPs are available, the RNA 3′ end is preferentially engaged by MUT-2, limiting RDE-8 access to the substrate (**Figure 5D**). *In vivo*, rNTPs are constitutively present, yet CPUG substrates are mRNAs that do not have an internal 3′ hydroxyl until RDE-8 cleavage generates one. Our *in vitro* data therefore suggest that once the substrate is cleaved, the new 3′ end is rapidly bound by MUT-2 for poly(UG)ylation, and that, once tailing begins, the RNA is no longer available for RDE-8-mediated cleavage.

### The substrate 3′ end determines the efficiency and register of poly(UG)ylation

Our tailing assays showed that ATP and CTP are poor substrates, whereas UTP and GTP support processive tail synthesis, consistent with MUT-2 acting as a poly(UG) polymerase as described previously^11–14^. These experiments, however, do not reveal whether tails generated by CPUG consist of strictly alternating UG repeats, nor whether the identity of the 3′-terminal nucleotides affects tailing efficiency.

To address this systematically, we incubated CPUG^+^ with a 37-nt single-stranded RNA pool bearing four randomized 3′-terminal nucleotides (4N RNA; 256 unique substrates; **Figure 6A**) in the presence of Mg^2+^ and all four rNTPs at equal concentrations (**Figure 6B** and **S6A**). Samples from 0 and 5 min time points were subjected to high-throughput sequencing (**Figure S6B**; **Table S5**). While CPUG^+^ generated long tails as seen by denaturing PAGE (**Figure S6A**), most sequencing reads contained tails of only 1-10 nucleotides, likely reflecting biases introduced during library preparation (**Figure 6C**). Across the substrate pool, incorporated nucleotides comprised 51.02% G and 48.79% U with negligible non-canonical incorporation (A, 0.01%; C, 0.18%) (**Figure 6C**). Position-resolved analysis of tail composition (n = 259,931 tails) revealed that at every tail position, G and U each accounted for ∼50% of incorporated nucleotides, and >97% of all dinucleotide steps corresponded to canonical GU or UG transitions (**Figure 6D**). Stratifying reads by U (n = 126,543 tails) or G (n = 131,574 tails) at position +1 revealed strictly alternating (UG)_n_ and (GU)_n_ tails, respectively (**Figure 6E**), demonstrating that the CPUG^+^ complex maintains strict UG alternation along the entire tail.

**Figure 6.**
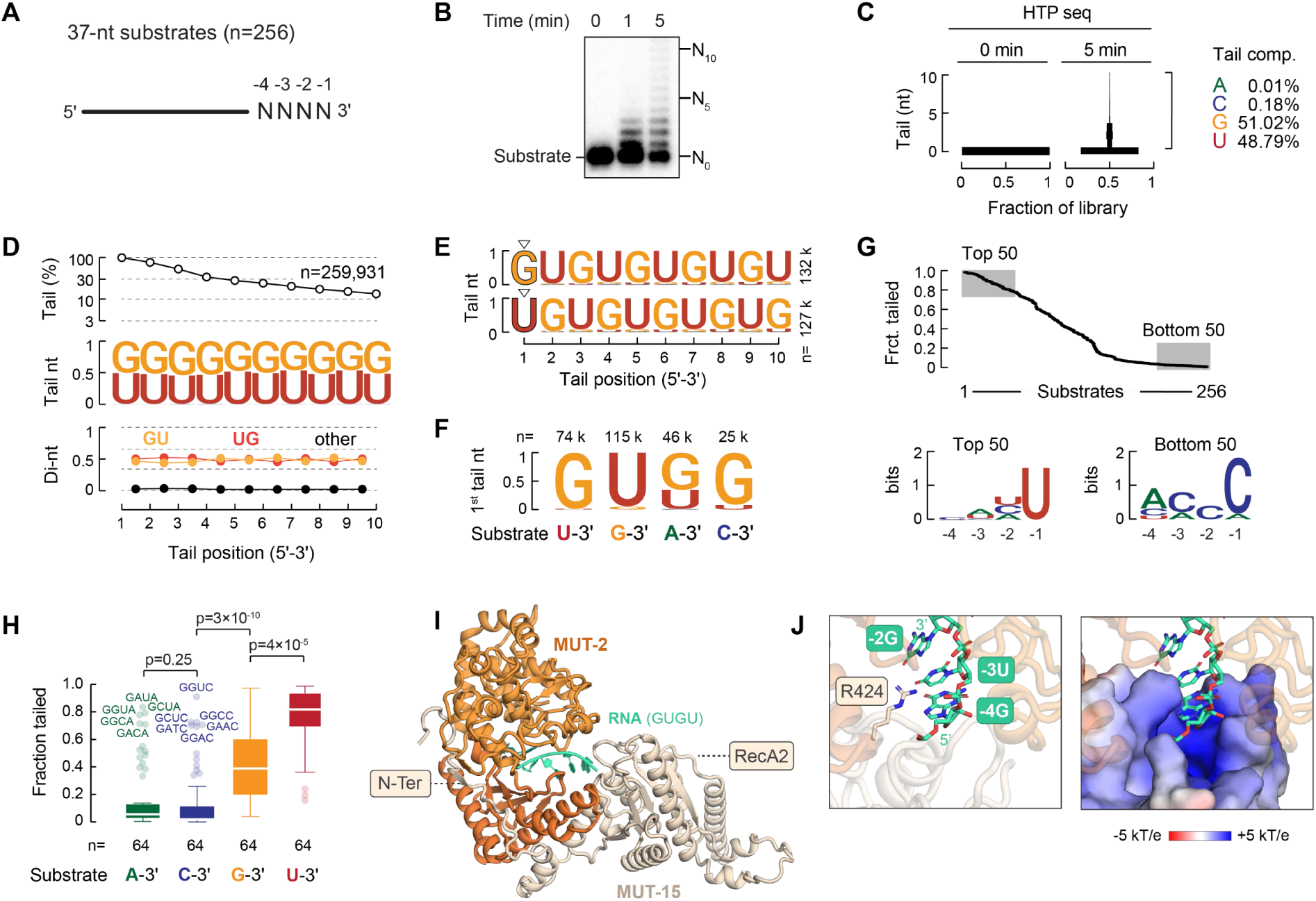
The substrate 3′ end determines the efficiency and register of poly(UG)ylation. **(A)** Schematic of the 5′-radiolabeled 37-nt 4N RNA pool used for high-throughput tailing assays. **(B)** Time-course tailing assay with CPUG^+^ and all four rNTPs. Positions of untailed substrate (N₀) and tailed species at +5 nt (N₅) and +10 nt (N₁₀) are indicated. The gel shows one of two independent replicates used for sequencing and analysis. **(C)** Tail-length distribution (0-10 nt) at 0 and 5 min determined by high-throughput sequencing. Right, nucleotide composition of all tails at 5 min. **(D)** Position-resolved tail composition of all tailed reads (n = 259,931 tails). Top, fraction of tails extending to each tail position. Middle, nucleotide composition at each tail position; Bottom, dinucleotide-step composition between adjacent tail positions. **(E)** Position-resolved tail composition for tails initiating with G (top; n = 131,574) versus tails initiating with U (bottom; n = 126,543). **(F)** Identity of the first incorporated nucleotide stratified by the 3′-terminal nucleotide of the substrate. Read numbers are indicated. **(G)** Tailing efficiencies (fraction tailed) of the 256 substrates. Sequence logos show positional information content (in bits) at each variable position (-4 to -1) for the 50 most and 50 least efficiently tailed substrates. **(H)** Tailing efficiencies grouped by 3′-terminal nucleotide (p-1) identity. Tukey box plots show median and interquartile range. Within the 3′-A and 3′-C classes, the most efficiently tailed members are labeled by their NNNN sequence. Adjusted p-values were calculated by the Kruskal-Wallis test with Dunn’s multiple-comparison correction. **(I)** AlphaFold3 prediction of the MUT-2/MUT-15 complex in the presence of a GUGU RNA substrate, GTP, and two Mg²⁺ ions. For MUT-15, only the N-terminal region and RecA2-like domain are shown. **(J)** Zoom-in from (I), with MUT-15 R425 highlighted. Right, electrostatic surface representation of the RecA2-like domain. See also Figure S6 and Table S5.

To determine whether the substrate 3′ end influences tailing, we analyzed tailing relative to the terminal nucleotide. Substrates ending in any of the four nucleotides were tailed, and the 3′-terminal nucleotide specified the register of poly(UG) synthesis: 3′-U and 3′-C substrates initiated with G, 3′-G substrates with U, and 3′-A substrates with either G or U (**Figure 6F**). Tailing efficiencies (fraction tailed) ranged from <1% to >99% across substrates. Sequence-logo analysis of the 50 most and least efficiently tailed substrates revealed a bias at the 3′-terminal (-1) position, with 3′-U enriched among efficiently tailed and 3′-C among poorly tailed RNAs (**Figure 6G**). Grouping by -1 identity confirmed a hierarchy: 3′-U substrates were tailed most efficiently (median 0.82), followed by 3′-G (0.39), whereas for 3′-A and 3′-C substrates the fraction tailed was much lower (0.06 and 0.02; **Figure 6H**). Unexpectedly, a subset of 3′-A and 3′-C substrates with a guanosine at the -4 position escaped this hierarchy, reaching efficiencies of 0.5-1.0, comparable to 3′-U substrates (**Figure 6H**). A heat map of all 256 substrates revealed pronounced sequence-dependent variation in tailing efficiency as a function of the randomized 3′ sequence (**Figure S6C**). Sorting substrates by -1 identity and then by G versus non-G at position -4 confirmed that a G at -4 raised efficiency for 3′-A and 3′-C substrates, showing that upstream sequence context modulates the dominant effect of the 3′-terminal nucleotide (**Figure S6D**).

The preference for guanosine at position -4 could arise from MUT-2 itself or from another CPUG component involved in RNA recognition. To explore a possible structural basis, we used AlphaFold3 to predict the MUT-2/MUT-15 complex together with RNA, Mg^2+^, and an incoming nucleotide. In the model, the disordered N-terminal region and the RecA2-like domain of MUT-15 both contact MUT-2 (**Figure 6I**), an interface that is conserved across *Caenorhabditis* species, including *C. remanei* and the distantly related *C. angaria* (**Figure S6E**). The positively charged MUT-15 RecA2-like domain lies adjacent to the catalytic cleft of MUT-2, with R425 near position -4 of the substrate RNA (**Figure 6I,J**). This arrangement suggests that MUT-15 extends the substrate-binding surface of MUT-2 and contributes to recognition of the upstream sequence context that influences tailing efficiency.

Together, our results define the CPUG complex as a register-locked alternating polymerase whose activity and register of poly(UG) synthesis are specified by the identity of the 3′-terminal substrate nucleotide, with additional modulation by upstream sequence context.

## Discussion

Small-RNA amplification in *C. elegans* is initiated by the conversion of Argonaute-recognized target transcripts into templates for RNA-dependent RNA polymerase (RdRP)-mediated 22G-RNA synthesis^4^. The CPUG complex mediates this conversion by cleaving and poly(UG)ylating the target RNAs ^9,12,13,16^. The present work provides a mechanistic framework explaining how these two enzymatic reactions are controlled and coordinated. Autoinhibition of the isolated RDE-8/NYN-1 nuclease module, together with its activation upon CPUG complex assembly, restricts cleavage to the appropriate molecular context. Moreover, the reduced accessibility of the RNA to RDE-8 while it is engaged by MUT-2, together with the resistance of the poly(UG) tail to cleavage, protects the RNA intermediate from re-cleavage and directs it toward RdRP-dependent 22G-RNA synthesis.

RDE-8 belongs to the Regnase family of nucleases, which has important functions in RNA regulation and cellular defense across animals^10,28,29^. In *C. elegans*, multiple family members have lost essential catalytic residues, including the pseudonuclease paralogs NYN-1 and NYN-2^10^. Whereas structurally characterized mammalian Regnase proteins form homodimers^28,30^, RDE-8 forms a stable heterodimer with NYN-1/2. The isolated RDE-8/NYN-1 complex is inactive, and its crystal structure reveals that the RDE-8 catalytic residues adopt an inactive conformation incompatible with metal-ion binding. Importantly, NYN-1 is not merely a passive dimerization partner: it contributes directly to the inhibitory network, helps form the catalytic cleft, and provides the main MUT-15-binding surface, thereby incorporating the nuclease module into CPUG.

Integration of RDE-8/NYN-1 into CPUG results in robust cleavage activity. Central to this activation is MUT-15, which we identify as a DEAD-box-like pseudohelicase. Its repurposed architecture bridges NYN-1 and MUT-2, linking the nuclease and tailing modules and potentially restricting cleavage to the fully assembled CPUG. MUT-15 binding, however, does not stabilize RDE-8 in a constitutively active conformation: in our cryo-EM structure, MUT-15 remodels the nuclease module, but the RDE-8 catalytic site remains inactive. Full activation may require substrate and metal-ion binding, with the RNA-binding activity of MUT-15 contributing to substrate recruitment or positioning. Yet MUT-15-mediated RNA recruitment cannot be the sole factor that drives activation, because the S159R mutation restores nuclease activity in the isolated RDE-8/NYN-1 complex, even in the absence of MUT-15. We therefore propose that MUT-15 acts as a licensing factor that allows RDE-8 to access catalytically competent conformations, with RNA and metal-ion binding completing active-site formation. A similar induced-fit mechanism, in which nonproductive catalytic aspartates are reorganized into a metal-bound site by partner proteins and RNA, has been proposed for the human NYN-domain nuclease MRPP3, the endonucleolytic subunit of the mitochondrial RNase P complex^42,43^. Such control would keep cleavage tightly restricted while ensuring efficient coupling to downstream processing.

Within CPUG, MUT-15 may also regulate MUT-2. MUT-15’s N-terminal region, which is necessary for MUT-2 binding *in vivo*^16^, wraps around a conserved hydrophobic surface on MUT-2 (**Figure S6F-H**). This surface is equivalent to the one of the *C. elegans* nucleotidyltransferase GLD-2 that is bound by its regulatory partners GLD-3 and RNP-8^44,45^ (**Figure S6I**). GLD-2 promotes translational activation of specific mRNAs during germline development, and these partners determine its activity and specificity by stabilizing the polymerase and stimulating polyadenylation^44,45^. MUT-15 may perform a similar role for MUT-2. Moreover, the MUT-15 RecA2-like domain lies near the MUT-2 catalytic cleft and could contribute to RNA positioning. To date, *in vitro* poly(UG) tailing has been reported only for the isolated *C. briggsae* MUT-2^14^, which produces shorter tails than CPUG. Whether this difference results from stimulation of MUT-2 within CPUG will require a comparison under identical experimental conditions.

But how does CPUG prevent re-cleavage of the RNA template or of the newly generated pUG tail? Our data identify two mechanisms that could impose directionality. First, CPUG appears to adopt mutually exclusive cleavage- and tailing-engaged states. *In vivo*, the cleavage-engaged state must occur first, because the target RNA initially lacks an internal 3′-OH. Binding of MUT-2 to the newly generated 3′ end would then promote the transition to the tailing-engaged state, in which the RNA is inaccessible to RDE-8. Re-cleavage would require the RNA to reposition and present a new internal segment. Second, RDE-8 does not efficiently cleave GU-alternating RNA, preventing poly(UG) tail cleavage. As the tested substrate is too short to form the characteristic pUG fold, this resistance is encoded by the alternating sequence itself.

Collectively, our findings support a model in which CPUG couples regulated RNA cleavage to poly(UG)ylation through multiple layers of control, converting a potentially hazardous endoribonucleolytic event into a committed entry point for small-RNA amplification.

A notable feature of CPUG is its architectural resemblance to the canonical cleavage and polyadenylation (CPA) machinery^46^. Both systems couple an active nuclease and an inactive paralog to a terminal nucleotidyltransferase^47^ within a larger RNA-processing assembly. In both cases, nuclease activity is only triggered within the fully assembled machinery^48,49^, allowing an irreversible cleavage event to be controlled and directly coupled to modification of the resulting RNA 3′ end which specifies the RNA’s subsequent fate. Although their components are distinct, the two systems illustrate a shared strategy for controlling catalysis and coordinating sequential RNA-processing reactions.

### Limitations of the study

Our reconstitution approach enabled direct analysis of CPUG architecture and enzymatic activities. However, the use of recombinant proteins and short non-physiological RNA substrates does not fully recapitulate the cellular context in which CPUG operates. Additional factors may regulate CPUG activity *in vivo*. Only part of CPUG was resolved by cryo-EM, so interactions involving MUT-2 and MUT-16 are inferred from AlphaFold models and crosslinking data. Moreover, none of our structures capture the RDE-8 catalytic site in an active conformation, so its activation and the transition to the tailing-engaged state remain models awaiting confirmation by RNA-bound structures. Finally, how Argonaute-recognized target RNAs are delivered to CPUG for RDE-8 cleavage remains unknown.

## Resource availability

### Lead contact

Requests for further information and resources should be directed to and will be fulfilled by the lead contact, Sebastian Falk.

### Materials availability

All unique reagents generated in this study are available from the lead contact without restriction.

### Data and code availability

#### Section 1: Data

- **Structures.** The atomic models have been deposited in the Protein Data Bank (PDB) under accession codes 28RS (RDE-8/NYN-1 wild-type complex, X-ray), 28RV (RDE-8/NYN-1 Ser159Arg mutant complex, X-ray) and 32VT (RDE-8/NYN-1/MUT-15 complex, cryo-EM). The cryo-EM map generated in this study has been deposited in the Electron Microscopy Data Bank (EMDB) under accession code EMD-59208. These data are publicly available as of the date of publication.
- **Mass spectrometry data.** The mass spectrometry proteomics data have been deposited to the ProteomeXchange Consortium via the PRIDE partner repository with the data set identifier PXD080438. These data are publicly available as of the date of publication.
- **Sequencing data.** Raw sequencing data will be deposited at GEO before publication and are available on request. Processed sequencing data are accessible in **Table S5**.
- **MD simulations.** Initial and final structures have been deposited on Zenodo (doi: 10.5281/zenodo.21836939) together with the processed trajectories. These data are publicly available as of the date of publication.

#### Section 2: Code

- **MD simulations.** All original codes have been deposited on Zenodo (doi: 10.5281/zenodo.21836939). These data are publicly available as of the date of publication.

#### Section 3: Additional information

Any additional information required to reanalyze the data reported in this paper is available from the lead contact upon request.

## Acknowledgements

We thank members of the Falk laboratory and Shamitha Govind for helpful discussions and support. The cryo-EM data collection was performed by the Electron Microscopy Facility at Vienna BioCenter Core Facilities (VBCF), member of the Vienna BioCenter (VBC), Austria. We thank Thomas Heuser and Harald Kotisch for assistance with data collection; Ingmar Schaefer, Thomas Lloyd Williams and Lorenz Emanuel Grundmann for suggestions on Cryo-EM data processing. AlphaFold predictions and Cryo-EM data processing were performed using the Life Science Compute Cluster (LiSC) of the University of Vienna. We acknowledge the European Synchrotron Radiation Facility (ESRF) for provision of synchrotron radiation facilities under proposal IDs MX2455 and MX2651 on beamline ID32-2. We thank Max Nanao for assistance and support during the beamtime. Proteomics analyses were performed at the Mass Spectrometry Facility of Max Perutz Labs using the VBCF instrument pool. Sequencing services were provided by the VBCF NGS Unit. We thank Thomas R. Burkard for processing the raw sequencing files. Molecular dynamics simulations were carried out using the Austrian Scientific Computing (ASC) infrastructure. We thank Bojan Zagrovic and Anton Polyansky for their input on MD simulations. This research was funded in whole or in part by the Austrian Science Fund (FWF) [10.55776/I6110, 10.55776/PAT4826925, 10.55776/ESP3222224, 10.55776/DOC177]. For open access purposes, the author has applied a CC BY public copyright license to any author-accepted manuscript version arising from this submission.

## Author contributions

V.B. and S.F. conceived the study. V.B. designed, performed and analysed experiments, and drafted the initial manuscript. S.F. contributed to the design and analysis of experiments. V.B. prepared samples for cryo-EM and protein crystallography. M.B. carried out cryo-EM data analysis and determined the cryo-EM structure with assistance from S.F. for model building. S.F. processed the X-ray diffraction data and solved the crystal structures. V.B. performed radioactive tailing assays and library preparation, with assistance from A.S. A.S. analysed sequencing data. M.M. performed MD simulations. V.B. and E.B. purified proteins. S.A. assisted with data analysis and interpretation. V.B. and S.F. prepared the final manuscript with input from all authors.

## Declaration of generative ai and ai-assisted technologies in the writing process

During the preparation of this work, the authors used ChatGPT (OpenAI) and Claude (Anthropic AI) to assist with language editing of the manuscript. After using these tools, the authors reviewed and edited the content as needed and take full responsibility for the content of the publication.

## Supplemental information

- **Document S1.** Supplemental Figures S1-S6, Tables S1-S4, Table S5 Legend, Movie S1 Legend, and Supplemental Methods
- **Table S5:** Processed sequencing data from CPUG RNA tailing assays
- **Movie S1:** MUT-15 binding is associated with remodeling of the nuclease module

## Material and Methods

### Recombinant protein expression and purification

Table S1 provides an overview of the constructs used in this study and the purification steps.

### Bacterial protein expression and purification

The genes coding for RDE-8 (Q23342) and NYN-1 (O18125) as well as the respective truncations were cloned into modified pET vectors. Proteins were produced as fusion constructs with an N-Ter His10 followed by different fusion tags in the *E. coli* BL21(DE3) or LOBSTER derivatives strain in Terrific Broth (TB) medium (**Table S1**). To reconstitute FL and truncated RDE-8/NYN-1 complexes, RDE-8 was co-expressed with NYN-1. Protein production was induced at 18°C by adding 0.2 mM IPTG for 12-16 hours. Cell pellets were resuspended in lysis buffer containing 50 mM NaH_2_PO_4_ pH 8, 20 mM TRIS/HCl pH 7.5, 250 mM NaCl, 20 mM imidazole, 10% glycerol, and 5 mM β-mercaptoethanol, supplemented with 0.01 mg/mL DNase, 1 mM PMSF and Benzonase (75U/mL), and cells were lysed by sonication. Clarified lysates were applied to a HisTrap FF column (Cytiva). After extensive washing with Lysis buffer and a chaperone wash (20 mM TRIS-HCl pH 7.5, 50 mM KCl, 10 mM MgCl_2_, 20 mM Imidazole, 10 % Glycerol, 5 mM ß-mercaptoethanol, 2 mM ATP), proteins were eluted with 500 mM imidazole in lysis buffer. Samples were dialyzed overnight at 4 °C in a buffer containing 20 mM TRIS/HCl pH 7.5, 150 mM NaCl, 10% glycerol, 20 mM imidazole, and 5 mM β-mercaptoethanol in the presence of 3C protease to remove affinity tags. Following tag cleavage, a His-tag chromatography was used to remove free tags. The flow-through was further purified by either anion-exchange chromatography using a linear NaCl gradient from 150 mM to 1 M NaCl (full-length RDE-8 and RDE-8/NYN-1) or heparin affinity chromatography using a linear NaCl gradient from 100 mM to 1 M NaCl (truncated RDE-8/NYN-1) (**Table S1**). Final purification was achieved by size-exclusion chromatography on a Superdex 200 Increase 16/600 column equilibrated in buffer containing 20 mM HEPES/NaOH pH 7.5, 150 mM NaCl, and 2 mM DTT. For proteins used for crystallization, DTT was replaced with 0.5 mM TCEP.

### Insect-cell protein expression and purification

Multigene expression constructs (RDE-8/NYN-1, MUT-15^52–541/MUT–1691–374^, RDE-8/NYN-1/MUT-15^52–541/MUT–1691–374^ and RDE-8/NYN-1/MUT-15^52–541/MUT–1691–374/MUT–2)^ were generated using a biGBac-based cloning strategy, and bacmid generation and baculovirus production were performed following the published biGBac protocol^50^. The genes coding for RDE-8 (Q23342) and NYN-1 (O18125) MUT-15 (Q22061), MUT-16 (O62011) and MUT-2 (O44768) fused to the desired Tags of interest (**Table S1**) were assembled into pLIB vectors by Gibson assembly. Codon-optimized MUT-15 cDNA was synthesized by Twist Bioscience. Gene expression cassettes were hierarchically assembled into the pBIG1a vector, and all constructs were verified by whole-plasmid sequencing. Recombinant bacmids were generated in DH10 EmBacY cells. Bacmids were transfected into Sf9 cells maintained in ESF 921 insect cell culture medium at 27 °C to generate recombinant baculovirus. 1L High Five cells, also maintained in ESF 921 medium at 27 °C, was infected with amplified baculovirus for protein expression and incubated at 21 °C for 5 days prior to harvesting by centrifugation. Pellets were resuspended in 50mL of desired lysis buffer supplemented with 0.01 mg/mL DNase, 1 mM PMSF and one complete Protease Inhibitor Cocktail tablet (Roche). Cell suspensions were stirred to homogenize and lysed by mild sonication. Benzonase (75U/mL) was added, and lysates were incubated for 30 min at 4 °C with stirring to reduce nucleic acid viscosity. Lysates were clarified by centrifugation prior to chromatography.

For recombinant RDE-8/NYN-1, RDE-8/NYN-1/MUT-15^52–541/MUT–1691–374^, RDE-8(D76N)/NYN-1/MUT-15^52–541/MUT–1691–374^ and RDE-8/NYN-1/MUT-15^52–541/MUT–1691–374/MUT–2^ complexes (Ce431, Ce421, Ce439 and Ce433, **Table S1**), pellets were resuspended in the following lysis buffer (50 mM NaH₂PO₄ pH 8.0, 20 mM TRIS-HCl pH 7.5, 250 mM NaCl, 10% glycerol, and 5 mM β-mercaptoethanol) and the N-terminal StrepII tag on RDE-8 was used for the initial affinity purification step. Clarified lysates were loaded onto a StrepTrap XT column (Cytiva). The column was washed extensively, and bound proteins were eluted using buffer supplemented with 50 mM biotin. Eluted complexes were dialyzed overnight at 4 °C in a buffer containing 20 mM TRIS/HCl (pH 7.5), 150 mM NaCl, 10% glycerol, and 5 mM β-mercaptoethanol in the presence of 3C protease to remove affinity tags. Following tag cleavage, the sample was applied to a heparin affinity column. Proteins were eluted using a linear NaCl gradient from 150 mM to 1 M NaCl. Peak fractions were pooled and concentrated using a 30 kDa molecular-weight–cutoff concentrator. Final purification was performed by size-exclusion chromatography on a Superdex 200 16/600 column equilibrated in buffer containing 20 mM HEPES/NaOH pH 7.5, 150 mM NaCl, 10% glycerol, and 2 mM DTT. Fractions corresponding to the desired complexes were pooled, concentrated, and stored at -70 °C for downstream analyses. For purification of the RDE-8/NYN-1 complex (Ce431), the same strategy was used, with the exception that a Q anion-exchange column (Cytiva) was employed in place of the heparin affinity column.

For MUT-15^52–541/MUT–1691–374^, pellets were resuspended in lysis buffer containing 50 mM NaH_2_PO_4_ pH 8, 20 mM TRIS/HCl pH 7.5, 250 mM NaCl, 20 mM imidazole, 10% glycerol, and 5 mM β-mercaptoethanol. Clarified lysates were applied to a HisTrap FF column (Cytiva). After extensive washing with Lysis buffer and a high-salt wash (50 mM NaH₂PO₄ pH 8.0, 20 mM TRIS-HCl pH 7.5, 500 mM NaCl, 20 mM imidazole, 10% glycerol, and 5 mM β-mercaptoethanol), proteins were eluted with elution buffer (50 mM NaH₂PO₄ pH 8.0, 20 mM TRIS-HCl pH 7.5, 250 mM NaCl, 500 mM imidazole, 10% glycerol, and 5 mM β-mercaptoethanol). Samples were dialyzed overnight at 4 °C in a buffer containing 20 Mm TRIS/HCl pH 7.5, 150 mM NaCl, 10% glycerol, 20 mM imidazole, and 5 mM β-mercaptoethanol in the presence of 3C protease to remove affinity tags. Following tag cleavage, the sample was further purified by heparin affinity chromatography using a linear NaCl gradient from 150 mM to 1 M NaCl. Final purification was achieved by size-exclusion chromatography on a Superdex 200 Increase 16/600 column equilibrated in buffer containing 20 mM HEPES/NaOH pH 7.5, 150 mM NaCl, 10% glycerol, and 2 mM DTT (**Table S1**).

### Mass spectrometry characterization of CPUG

Recombinant CPUG^+^ (Ce433; **Table S1**; **Figure 1E**) was characterized by intact, bottom-up, and cross-linking mass spectrometry to assess its molecular mass, protein composition, and inter-subunit contacts. Detailed methods are provided in the Supplementary Methods.

### X**-** ray crystallography

#### Sample preparation and crystallization

Truncated constructs of *C. elegans* RDE-8 (residues 1-300) and NYN-1 (residues 115-500) were designed based on AlphaFold2 predictions and co-expressed as an MBP-tagged complex in *E. coli* (**Table S1**). For crystallization trials, purified complexes were diluted in storage buffer (20 mM HEPES/NaOH pH 7.5, 150 mM NaCl, and 0.5 mM TCEP) to a final concentration of 4 mg/ml. Crystallization experiments were set up using hanging-drop vapor diffusion in 24-well plates, with drops consisting of protein and reservoir solution mixed at ratios of 1:1 or 1:2, and incubated at 4 °C. Crystals of RDE-8/NYN-1 complexes reproducibly formed in conditions containing PEG 3350 as precipitant (typically 13-21%) and 200 mM KSCN. For crystallization attempts in the presence of divalent cations, MgCl₂ or MnCl₂ was included at concentrations ranging from 2 to 20 mM. Crystals were harvested and cryoprotected by transfer into reservoir solution supplemented with increased PEG concentration (typically +2%) and glycerol (20%) and vitrified in liquid nitrogen prior to data collection.

#### Structure determination

Diffraction data for the RDE-8/NYN-1 wild-type complex and the RDE-8 (S159R mutant)/NYN-1 complex were collected at beamline ID23-2 of the European Synchrotron Radiation Facility (ESRF, Grenoble, France). Diffraction data of the RDE-8 / NYN-1 wild-type complex were processed automatically by the Grenoble Automatic Data Processing (GrenADES) pipeline at the ESRF^51^. Within this pipeline, data were integrated with XDS^52^ and further processed using POINTLESS and AIMLESS^53,54^. The diffraction data of the RDE-8 (S159R mutant) / NYN-1 complex were processed automatically by the xia2 pipeline at the ESRF. Within this pipeline, data were integrated with DIALS^55^ and scaled using AIMLESS^54^. Phases were determined by molecular replacement using the best AlphaFold3^36^ models of *C. elegans* RDE-8 (residues 1 to 300) and NYN-1 (residues 115 to 500). Prior to molecular replacement, the models were prepared with Phenix (process_predicted_model) to convert pLDDT values into B factors and to remove flexible regions. Molecular replacement was performed with Phaser^56^ within Phenix^57^. The resulting model was automatically built with ModelCraft^58^ within CCP4i2^59^, completed manually with Coot^60^ and refined with phenix.refine^61^ and REFMAC5^62^. The final models are well ordered and most of both chains was modeled with high confidence, while regions of lower confidence, mainly the mobile N-terminal segment of RDE-8 (residues 2 to 28) and surface loops in NYN-1 (122-123 and 128-129, 230 to 236 and 425 to 434), showed elevated B-factors and were modeled with support from the AlphaFold3 model. Model quality was assessed with MolProbity^63^ and PDB-REDO^64^. Data collection and refinement statistics are given in **Table S2**. Molecular graphics were generated with The PyMOL Molecular Graphics System, Version 3.0 Schrödinger, LLC.

### Molecular Dynamics Simulations

Atomistic molecular dynamics simulations were run with GROMACS 2024.4^65^. All simulations started from the experimentally determined X-ray structure (this study, PDB ID: 28RS). The S159R variant was generated by in silico substitution of the same structure, so the wild-type and mutant systems differ only by this residue. Additional information about simulations is available in (**Table S3**). The Amber99sb-ILDN force field^66^ and the TIP3P^67^ water model were used. Parameters for the divalent ion manganese were obtained from Bradbrook et al.^68^. All aspartate and glutamate residues were modelled in their deprotonated states, and histidine protonation was assigned by pdb2gmx on the basis of the local hydrogen-bonding network. The Mn^2+^ ion was positioned manually at the centroid of the catalytic aspartate cluster. Each system was placed in a truncated octahedron box with a minimum distance of 1.5 nm between the protein complex and the box edges, and subsequently solvated. The system was neutralized by the addition of counterions, no salt was added. The equations of motion were integrated with a 2 fs time step. Bonds involving hydrogen atoms were constrained using the LINCS algorithm^69,70^ (lincs_order = 4, lincs_iter = 1). Long-range electrostatics were described by the Particle Mesh Ewald (PME) method, with a grid spacing of 0.12 nm and a cut-off of 1.2 nm for real-space interactions. Van der Waals interactions were truncated at 1.2 nm using the Verlet cut-off scheme, and long-range dispersion corrections were applied to both energy and pressure. To maintain the temperature at 300 K the Bussi-Donadio-Parrinello thermostat^71^ was used, with a coupling constant τT of 0.1 ps. With the Parrinello-Rahman barostat^72^ we maintained the pressure at 1 bar, with a coupling constant τp = 2 ps. Energy minimization with the steepest descent algorithm was run for 50,000 steps or until the maximum force was less than 1000 kJ/mol/nm. The system was equilibrated in the NVT ensemble for 100 ps, followed by 100 ps of equilibration in the NPT ensemble. No position restraints were applied at any stage. Five independent replicates per variant were generated by assigning different randomly seeded initial velocities during NVT equilibration and production runs continued from the corresponding NPT checkpoints. Each replicate was run for 5 μs, with coordinates saved every 10 ps. Thermostat and barostat settings during production run were as in the NPT equilibration. Trajectories were analysed using the MDAnalysis and mdtraj Python packages ^73–75^, using the full production run.

### Cryo-EM Structure

#### Sample preparation

Purified CPUG^+^ complex was mixed with a 1.5-fold molar excess of a 28-nt RNA substrate in a buffer containing 20 mM HEPES/NaOH, 150 mM NaCl, and 2 mM DTT, and incubated on ice for 45 min. The RNA (GGUGCAUCUAAAGUUGAUUGAAGAGUUC) was synthesized *in vitro* as described in Pereirinha et al.^76^. The sample was subsequently cross-linked with 0.5 mM DSBU for 30 min at room temperature in a final reaction volume of 50 µL. The reaction was quenched by addition of glutamic acid monosodium salt pH 7.5 to a final concentration of 50 mM, followed by a further 15 min incubation at room temperature. Uncrosslinked control samples were prepared in parallel under identical conditions without addition of DSBU. The crosslinked sample was subjected to size-exclusion chromatography on a Superdex 200 increase 3.2/300 column using a microÄKTA system. This resulted in an A260/A280 ratio of approximately 0.6, compared with ∼1 for the uncrosslinked control, which remained stably associated with RNA. This decrease indicates that crosslinking displaced the bound RNA. Consequently, the cryo-EM structure represents the RNA-free state of the CPUG^+^ complex. Peak fractions corresponding to the assembled complex were pooled. Peak fractions were diluted to 1.5 µM in 20 mM HEPES/NaOH pH 7.5, 150 mM sodium chloride, 2 mM DTT and supplemented with 0.005% of fluorinated octyl-ß-maltoside. Quantifoil Cu 300 mesh R1.2/1.3 grids were glow-discharged before applying 4 µl sample. Grids were blotted for 2 seconds at 4 °C and 85 % humidity and plunged into liquid ethane for vitrification using a Leica EM GP. **Data collection and processing**

Data were collected using a 300 kV Titan Krios G4 (Thermo Fisher Scientific) equipped with a cold field emission gun, an energy filter with a slit width of 10 eV and a Falcon 4 direct electron detector. Movies were acquired in EER-format at a nominal magnification of 165 000-fold, corresponding to a pixel size of 0.749 Å/px, with a defocus range between -2.5 and -1.0 µm and a cumulative fluence of 40 e/Å^2^. All processing steps were performed using cryoSPARC (5.0.6)^77^. Following patch motion correction, patch CTF estimation and manual curation of 16,001 movies, 14,652 movies were used as input for initial particle picking using the blob picker in a range from 60 to 160 Å. Particles were extracted with a box size of 400 pixels and sorted iteratively by 2D classification and heterogenous refinement. High-quality 2D classes were used to train Topaz for particle picking^78^. Once extracted, particles were again sorted iteratively by 2D classification and heterogenous refinement. Non-uniform refinement followed by masked local refinement around RDE-8/NYN-1/MUT-15 yielded a high-resolution density that was used for reference-based motion correction^79^. A total of 151,515 particles were re-extracted, followed by global and local CTF refinement. Non-uniform refinement with a mask around RDE-8/NYN-1/MUT-15 resulted in a map with an overall resolution of 2.9 Å. Local resolution estimation was performed in cryoSPARC.

#### Model building and refinement

Initial coordinates were generated by docking AlphaFold3 predictions^36^ of RDE-8, NYN-1 and MUT-15 into the cryo-EM density using Phenix (version 2.0.5867)^57^. Models were iteratively refined through manual adjustment in Coot (version 0.9.8.95)^60^ and Isolde (version 1.10.1)^80^ and real-space refinement in Phenix^61^. Model building for RDE-8 and NYN-1 was additionally guided by the high-resolution crystal structures obtained within this study (PDB code: 28RS). Owing to weaker local density, several regions were modeled with lower confidence: MUT-15 residues 75 to 78 and 176 to 180, and NYN-1 residues 426 to 430. Model statistics were evaluated with Molprobity^63^ within Phenix and DAQscore^81^ and are summarized in Table S4. Graphical representations were prepared using PyMOL (version 3.1.3) and ChimeraX (version 1.10)^82^.

### RNA cleavage assays

Endoribonuclease assays were performed using single-stranded RNA substrates carrying a 5′ 6-fluorescein amidite (5′-FAM) label, purchased from Ella Biotech (Fuerstenfeldbruck, Germany). The following 16-nt RNAs were used: RNA-1 (5′-GUUGAUUGAAGAGUUC-3′), RNA-1(G_8_A) (5′-GUUGAUUAAAGAGUUC-3′), RNA-1(G_11_A) (5′-GUUGAUUGAAAAGUUC-3′), RNA-2 (5′-AGCACCGUAAAGACGC-3′), pA (5′-AAAAAAAAAAAAAAAA-3′), pU (5′-UUUUUUUUUUUUUUUU-3′), pUG (5′-GUGUGUGUGUGUGUGU), and pUG(U_10_A) (5′-GUGUGUGUGAGUGUGU-3′). Reactions were performed in 20 mM Tris-HCl (pH 7.5), 100 mM KCl, 2 mM DTT, 2 mM MnCl₂, and 2 mM MgCl₂, supplemented with RNasin (Promega) at 1 U/µL. For ion-dependency assays, reactions were performed either without added divalent cations, with 2 mM MgCl₂, or with 2 mM MnCl₂. RNA substrates were used at 500 nM and proteins or protein complexes at 1 µM. Protein complexes were preassembled on ice for 30 min before addition to the reaction. Reactions were incubated for 30 min at room temperature and terminated by addition of an equal volume of Gel Loading Buffer II (Thermo Fisher Scientific). Samples were heated at 95 °C for 5 min and resolved by 15% denaturing urea-PAGE (19:1 acrylamide:bis-acrylamide) in 1× TBE. Gels were pre-run at 250 V for 45 min and electrophoresed at 200 V for 45 min. FAM-labeled RNA products were visualized by fluorescence scanning using a Typhoon imager.

### RNA tailing assays

RNA tailing assays were performed using purified CPUG^+^ at a final concentration of 500 nM and 28-nt RNA substrates (3′OH-RNA, 5′-AUUGCAUCUAAAGUUGAUUGAAGAGUUC-3′; 3′ddC-RNA, 5′-AUUGCAUCUAAAGUUGAUUGAAGAGUUddC-3′) at 500 nM. The RNAs carried a 5′-6-fluorescein amidite (FAM) label and were purchased from Ella Biotech (Fuerstenfeldbruck, Germany). For nucleotide-specificity time-course experiments, reactions were performed in 20 mM Tris-HCl (pH 7.5), 100 mM KCl, 2 mM DTT, 20 mM MgCl₂, and 0.025% NP-40. ATP, CTP, GTP, or UTP was supplied individually at 2.5 mM; where UTP and GTP were combined, each nucleotide was present at 2.5 mM, and reactions containing all four rNTPs contained 2.5 mM of each rNTP. Reactions were performed in protein low-binding tubes (Eppendorf) in a final volume of 30 µL, and aliquots were withdrawn at different time points. For divalent-cation dependence assays, reactions were performed in 20 mM Tris-HCl (pH 7.5), 100 mM KCl, and 2 mM DTT, supplemented with either 5 mM MgCl₂ or 5 mM MnCl₂. ATP was added at 1.25 mM, whereas UTP and GTP were each added at 1.25 mM where indicated. For assays comparing 3′OH-RNA and 3′ddC-RNA, reactions were performed in 20 mM Tris-HCl (pH 7.5), 100 mM KCl, 2 mM DTT, and 5 mM MnCl₂. Where indicated, ATP was supplied at 1.25 mM, or UTP and GTP were each supplied at 1.25 mM. Reactions were incubated for 5 min. All reactions were terminated by addition of an equal volume of loading dye followed by heating. Reaction products were resolved on 15% denaturing polyacrylamide gels in 1× TBE. Gels were pre-run at 250 V, and samples were electrophoresed at 200 V. FAM-labeled RNA products were visualized by FAM fluorescence using a Typhoon scanner. For nucleotide-specificity time-course experiments, gels were subsequently stained with SYBR Gold (5 µL in 50 mL 1× TBE) for 5 min and imaged directly.

### High throughput sequencing

#### *In vitro* tailing assays

The synthetic RNA (ACACUCUUUCCCUACACGACGCUCUUCCGAUCUNNNN) was purchased from Horizon. For radiolabeling, 30 pmol RNA was 5′-end-labeled with γ-32P-ATP (6,000 Ci/mmol; Hartmann Analytic) using T4 polynucleotide kinase (PNK; NEB). Radiolabeled RNA was purified using G25 spin columns (GE Healthcare) and further resolved on 10% denaturing PAGE. *In vitro* tailing reactions were performed at 22 °C with 20 nM radiolabeled RNA and 500 nM recombinant CPUG^+^ complex in 10 µl reaction buffer (20 mM TRIS-HCl pH 7.5, 100 mM KCl, 2 mM DTT, 0.025% NP-40, 20 mM MgCl₂, and 2.5 mM of each rNTP) using protein low-binding tubes (Eppendorf). Reactions were initiated by addition of the CPUG^+^ complex. Samples collected at the indicated time points were subjected to acidic phenol-chloroform extraction followed by ethanol precipitation for sequencing experiments. A fraction of each extracted sample was analyzed by denaturing PAGE, while the remaining RNA was used for sequencing library preparation. Gels were dried and visualized using a Storm PhosphorImager (GE Healthcare).

#### RNA library preparation and sequencing analysis

Libraries for high-throughput RNA tailing were generated as previously described^83^. Briefly, RNA recovered from the in vitro RNA assays by phenol-chloroform extraction was ligated to 3′ adapter containing 4 random nucleotides at the ligation interface to minimize ligation bias. Reverse transcription was performed using SuperScript III Reverse Transcriptase (Invitrogen), followed by PCR amplification using NEBNext Ultra II Q5® Master Mix (NEB). Amplified cDNA products were purified on 2% agarose gels. Library quality control and high-throughput sequencing were performed by the VBCF Next Generation Sequencing facility. For high-throughput sequencing-based tailing analyses, reads per million (RPM) values corresponding to individual tail sequences were combined across two biological replicates. To visualize nucleotide preferences across the 4N region, substrates corresponding to the top 50 or bottom 50 most/least tailed were selected. Position probability matrices (PPMs) were computed from selected sequences. Information content (I^seq^) was calculated following Stormo et al. (2000) using the RPM-weighted input PPM as background^84^. Negative information content values (nucleotide depletion relative to input) were removed from plots. Logos were generated using ggseqlogo^85^ (v0.26; R).

Statistical analyses were performed in Prism v10.1.1 (GraphPad), Excel v16.80 (Microsoft), or R (v4.4.2; 2024-10-31 ucrt).

### Writing

ChatGPT (OpenAI) and Claude (Anthropic AI) were used to assist with language editing of the manuscript. All authors reviewed and take responsibility for the content.

