## Supplementary material for "A pseudohelicase-centered complex couples assembly-dependent RNA cleavage to poly(UG)ylation": Document S1

[1] Max Perutz Labs, Vienna BioCenter, 1030 Vienna, Austria
[2] University of Vienna, Vienna, Austria
[3] Vienna BioCenter PhD Program, a Doctoral School of the University of Vienna and the Medical University of Vienna, 1030 Vienna, Austria
[4] University of Vienna, Vienna Doctoral School in Chemistry (DoSChem), 1090 Vienna, Austria.
[5] Institute of Molecular Biology, Mainz, Germany
[6] PhD Program on Gene Regulation, Epigenetics & Genome Stability, Mainz, Germany
[7] Present address: Center for Physiology and Pharmacology, Institute of Pharmacology, Medical University of Vienna, 1090 Vienna, Austria
[8] Johannes Gutenberg University Mainz, Mainz, Germany
[9] Institute of Molecular Biotechnology, IMBA, Vienna BioCenter (VBC), 1030 Vienna, Austria
[10] These authors contributed equally to this work
[11] Lead contact

*Correspondence:
,


#### Supplemental contents:

- Figures S1-S6
- Tables S1-S4
- Table S5 Legend
- Movie S1 Legend
- Supplemental Methods
- Supplemental References

### Supplemental Figures

#### Figure S1. The CPUG complex is organized around a pseudohelicase

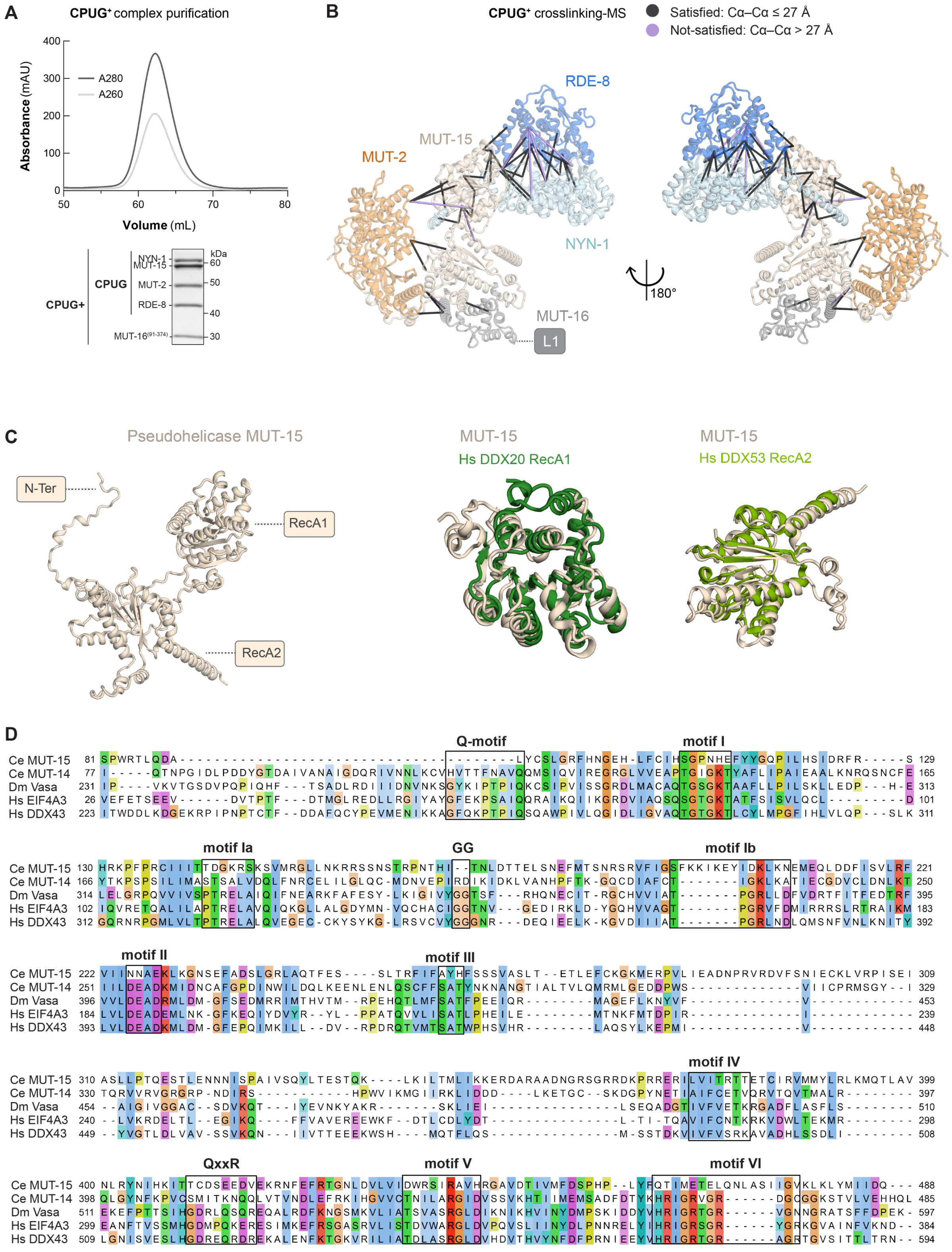

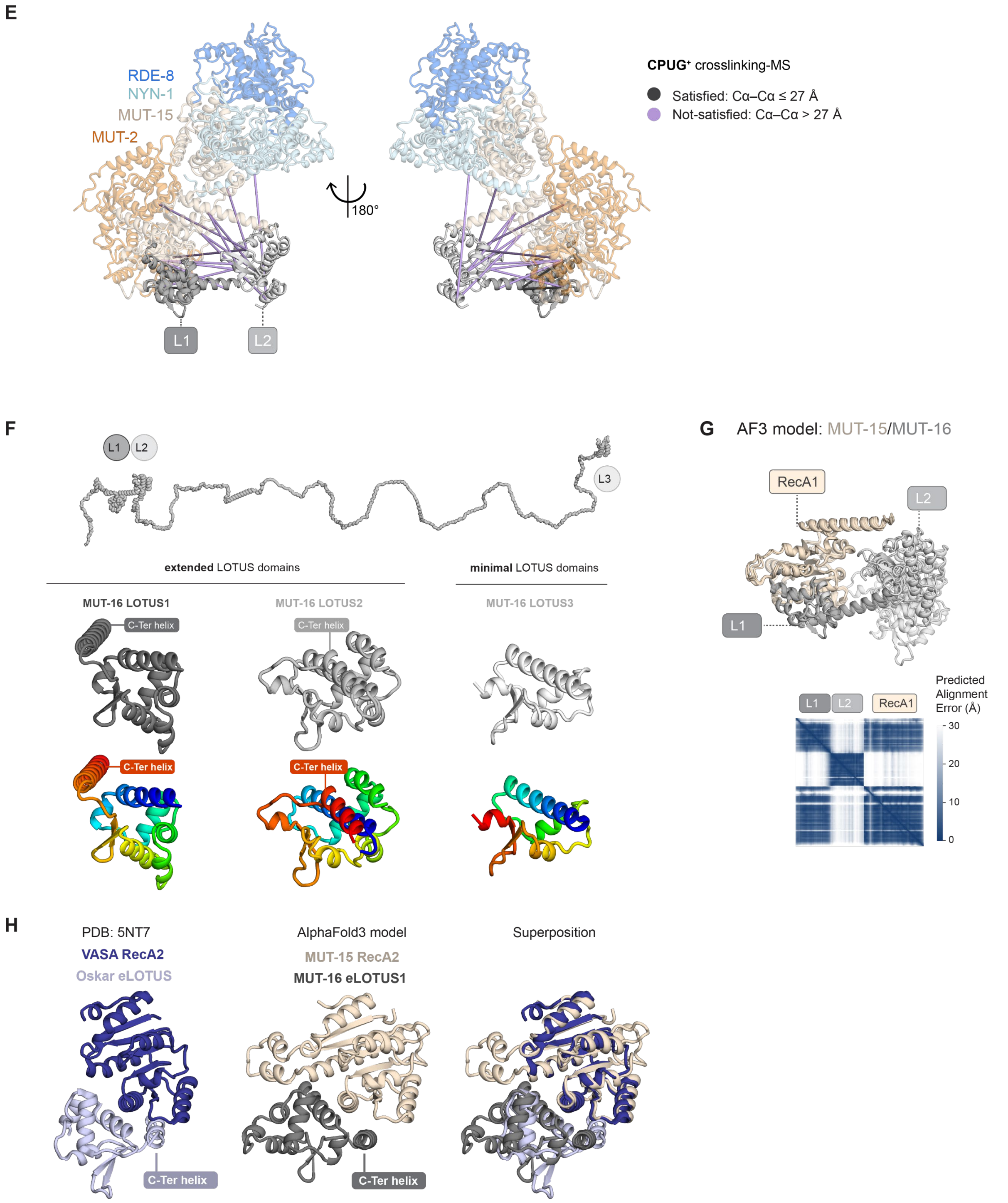

**(A)** Size-exclusion chromatography profile of the purified CPUG⁺ complex on a HiLoad 16/600 Superdex 200 column. The values of absorbance at 260 nm ($A_{260}$) and 280 nm ($A_{280}$) are shown. Peak fractions were analyzed by SDS–PAGE and Coomassie staining. The gel is also shown in Figure 1E. **(B)** Intermolecular crosslinks between CPUG⁺ components mapped onto the AlphaFold3 model of the complex. Distance-compatible crosslinks, defined by a Cα–Cα distance of ≤27 Å (Satisfied), are shown in dark gray; distance-incompatible crosslinks, defined by a Cα–Cα distance of >27 Å (Not-Satisfied), are shown in lavender. For clarity, only the first LOTUS domain of MUT-16, L1, is shown; crosslinks involving L2 are shown in (E). Two views of the model are presented. **(C)** AlphaFold3 model of MUT-15 and structural comparison of its two folded domains with their closest Foldseek matches. The N-terminal domain aligns with the RecA1 domain of human DDX20 (PDB: 2OXC; TM-score, 0.63; RMSD, 8.0 Å), whereas the C-terminal domain aligns with the RecA2 domain of human DDX53 (PDB: 8KCA; TM-score,

0.74; RMSD, 8.7 Å). **(D)** Sequence alignment of *C. elegans* MUT-15 and representative DEAD-box helicases. The alignment was generated using Clustal Omega and visualized in Jalview. Sequences correspond to *C. elegans* MUT-14 (UniProt: Q17978), *D. melanogaster* Vasa (UniProt: P09052), human eIF4A3 (UniProt: P38919), and human DDX43 (UniProt: Q9NXZ2). Conserved sequence motifs characteristic of DEAD-box helicases are highlighted[1,2]. **(E)** Intermolecular crosslinks between MUT-$16^{91–374}$ and the CPUG components mapped onto the AlphaFold3 model, as in (B). MUT-16 LOTUS1 (L1) and LOTUS2 (L2) domains are highlighted. **(F)** AlphaFold3 models of the three structured domains of MUT-16, identified here as LOTUS domains L1 (eLOTUS, residues 116-235), L2 (eLOTUS, residues 239-374), and L3 (mLOTUS, residues 974–1054). The three domains are shown in shades of grey or colored from the N-terminus (in blue) to the C-terminus (in red). **(G)** AlphaFold3 model of the MUT-15 RecA2-like domain bound to MUT-$16^{91-374}$. The L1 and L2 LOTUS domains of MUT-16 are shown in light and dark gray, respectively. The five generated models are superimposed to illustrate the flexibility of L2 relative to L1. The predicted aligned error (PAE) plot corresponds to the highest-ranked AlphaFold3 model and is representative of the five generated models. **(H)** AlphaFold3 model of the MUT-15 RecA2-like domain bound to MUT-16 L1 is shown alongside the *D. melanogaster* Vasa RecA2/Oskar LOTUS domain complex (PDB: 5NT7). A superposition of the two complexes is shown on the right.

### Figure S2. The nuclease module adopts an autoinhibited conformation

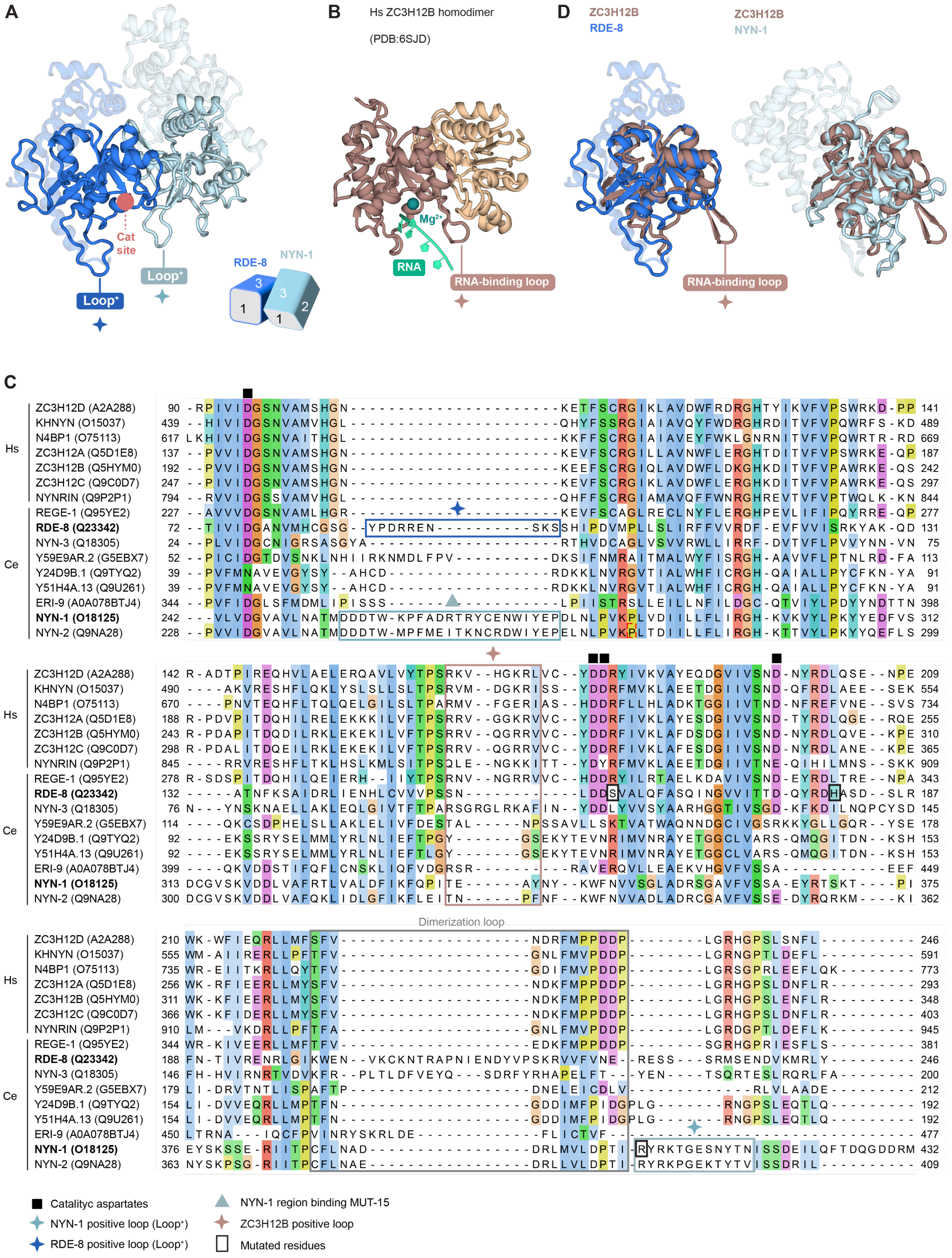

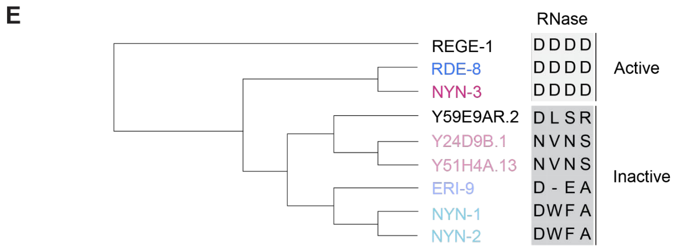

F

| | | Active | | Inactive | | | | | |
|---|---|---|---|---|---|---|---|---|---|
| | | RDE-8 | NYN-3 | Y59E9AR.2 | Y24D9B.1 | Y51H4A.13 | ERI-9 | NYN-1 | NYN-2 |
| Active | RDE-8 | 0.039 | 0 | 0 | 0 | 0.083 | 0.41 | 0.545 | 0.48 |
| | NYN-3 | 0 | 0.085 | 0 | 0.576 | 0.603 | 0 | 0 | 0 |

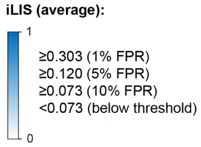

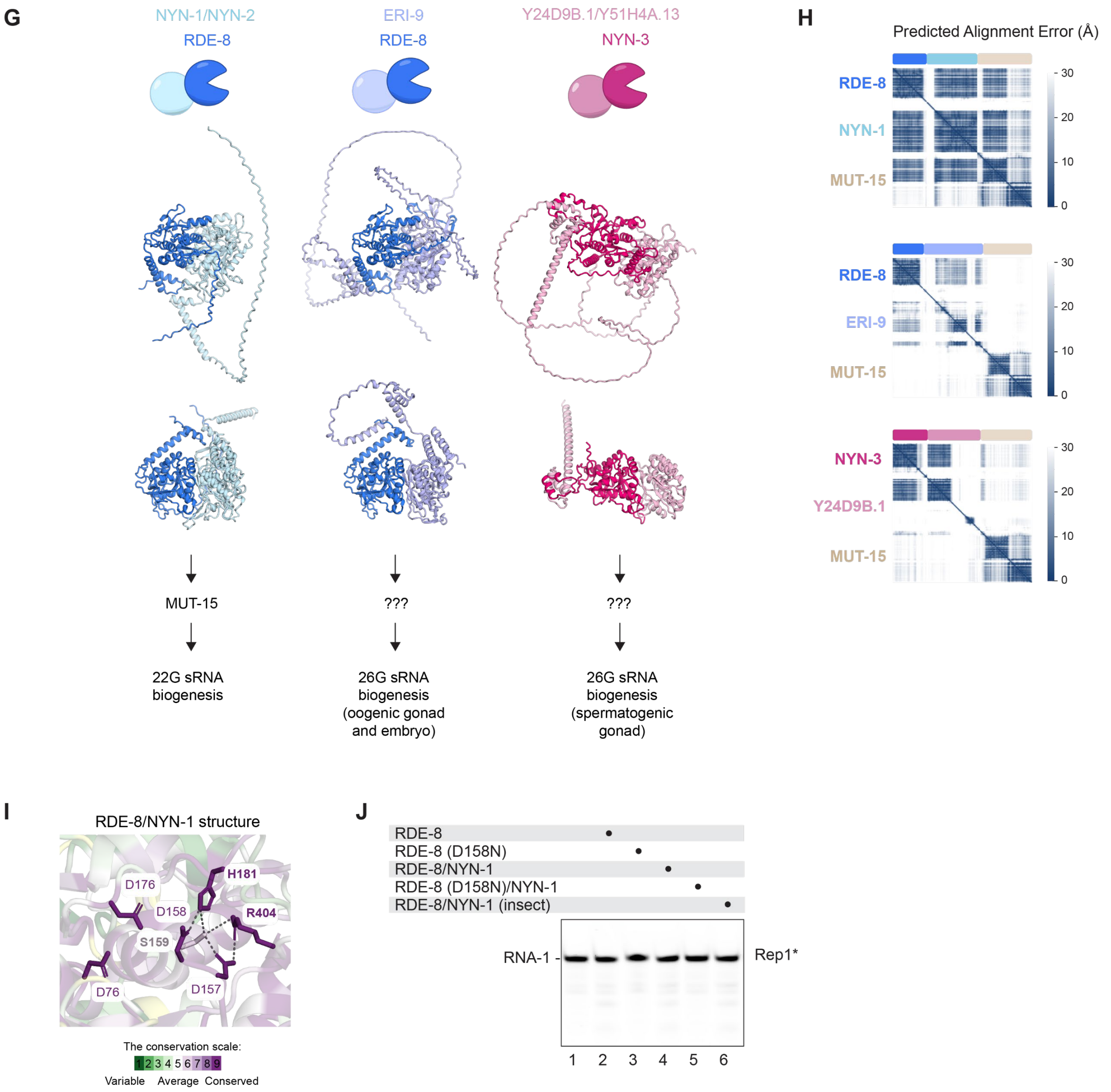

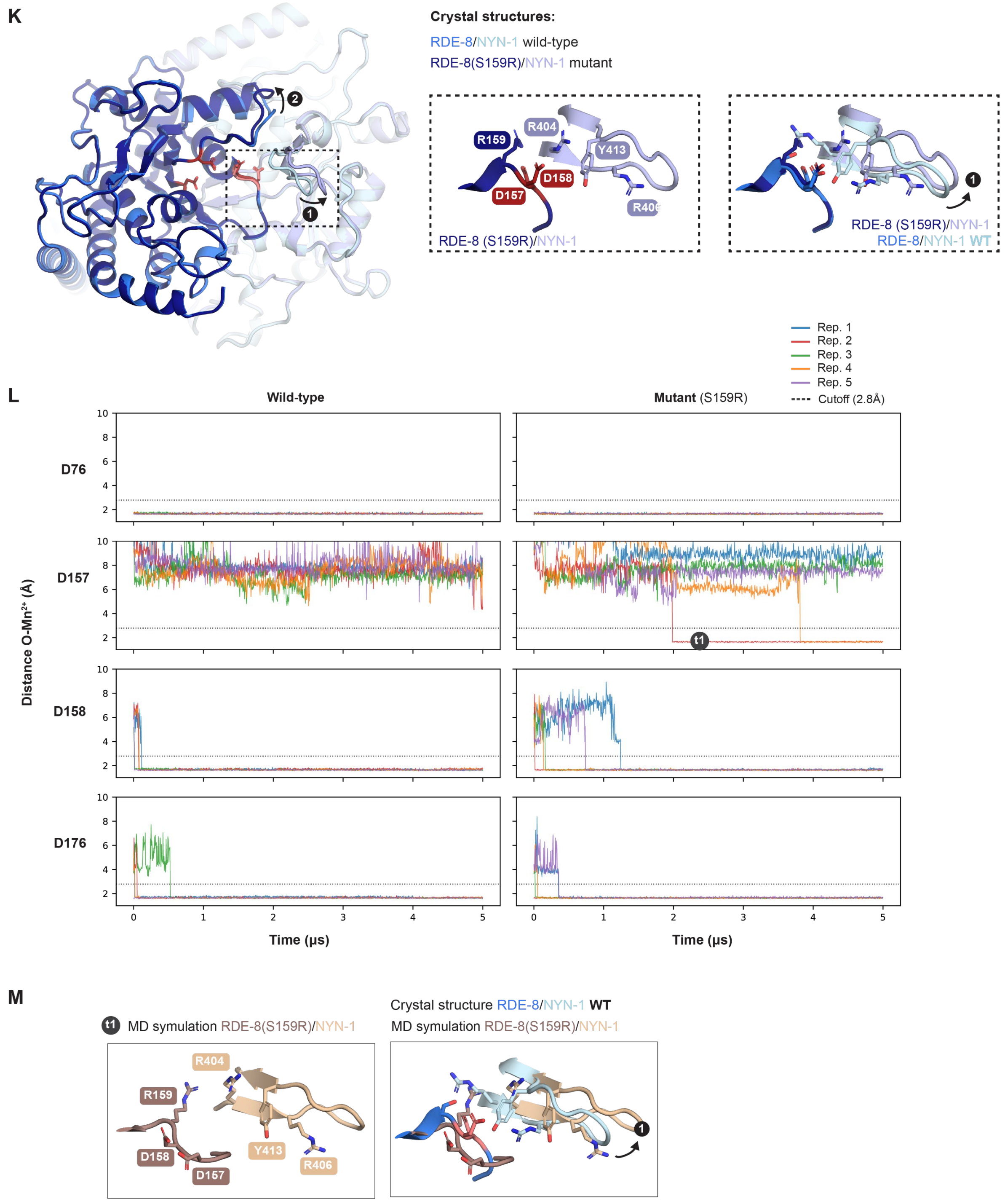

**(A)** Crystal structure of the *C. elegans* RDE-8/NYN-1 heterodimer (This study, PDB: 28RS). Regions extending beyond the minimal NYN domains are shown with transparency. The schematic illustrates the relative orientation of the RDE-8 and NYN-1 NYN domains. The NYN domains are tilted and asymmetric, with that of NYN-1 rotated by approximately 72° relative to that of RDE-8. The RDE-8- and NYN-1/2-specific positively charged insertions are indicated as in (C). **(B)** Crystal structure of human ZC3H12B bound to RNA (PDB: 6SJD). ZC3H12B RNA-binding loop is indicated as in (C). **(C)** Structure-based sequence alignment of human and *C. elegans* Regnase-family proteins. Sequences were selected as described by Habacher and Ciosk[3], and the alignment was generated with PROMALS3D using the AlphaFold3 models of the NYN domains. Conserved catalytic residues and the RNA-binding and dimerization loops described for mammalian Regnase proteins are indicated. NYN-1/2- and RDE-8-

specific insertions and residues mutated in this study (RDE-8 S159R, RDE-8 H181A and NYN-1 R404A) are highlighted. **(D)** Structural superposition of the NYN domains of RDE-8 (left) and NYN-1 (right) with the NYN domain of ZC3H12B (PDB: 6SJD). The ZC3H12B RNA-binding loop, which is absent in RDE-8 and NYN-1, is highlighted. **(E)** Phylogenetic tree of the *C. elegans* Regnase family. The presence of the conserved DDDD catalytic motif, indicative of an active RNase domain, or the corresponding substitutions in predicted inactive family members is indicated. Adapted from Habacher and Ciosk[3]. **(F)** AlphaFold3 predictions of pairwise interactions between the catalytically active Regnase-family proteins RDE-8 (UniProt: Q23342) or NYN-3 (UniProt: Q18305) and the indicated *C. elegans* Regnase-family members: RDE-8 (Q23342), NYN-3 (Q18305), Y59E9AR.2 (G5EBX7), Y24D9B.1 (Q9TYQ2), Y51H4A.13 (Q9U261), ERI-9 (A0A078BTJ4), NYN-1 (O18125), and NYN-2 (Q9NA28). Values represent the mean integrated Local Interaction Score (iLIS) calculated from five AlphaFold3 models using LIVIA (https://flyark.github.io/LIVIA/)[4]. **(G)** Cartoon representations of the AlphaFold3-predicted RDE-8/NYN-1, RDE-8/ERI-9, and NYN-3/Y24D9B.1 heterodimers, shown with (upper) or without (lower) predicted disordered regions. The scheme summarizes our model, in which catalytically active Regnase-family proteins associate with inactive members that contribute to their recruitment to distinct small-RNA pathways. **(H)** AlphaFold3 predictions of interactions between MUT-15 and the active/inactive couples: RDE-8/NYN-1, RDE-8/ERI-9, and NYN-3/Y24D9B.1. PAE plots for the highest-ranked model of each predicted complex are shown. **(I)** Close-up view of the RDE-8 active site in the RDE-8/NYN-1 crystal structure (this study, PDB: 28RS) highlighting residues involved in autoinhibition (RDE-8 S159 and H181, and NYN-1 R404) colored according to ConSurf[5] conservation scores. **(J)** RDE-8-mediated RNA cleavage was assessed using RDE-8 or the RDE-8/NYN-1 heterodimer, and a 16-nt 5′-FAM-labeled RNA substrate (RNA-1). RDE-8/NYN-1 was purified from bacteria or insect cells. Catalytically inactive RDE-8 and RDE-8/NYN-1 (D158N) were included as negative controls. Cleavage products were resolved by denaturing PAGE and detected by FAM fluorescence. Full gel and replicates are shown in Figure S3A. **(K)** Comparison of the crystal structure of the RDE-8(S159R)/NYN-1 heterodimer (this study, PDB: 28RV) with that of the wild-type RDE-8/NYN-1 complex (This study, PDB: 28RS). The four RDE-8 catalytic aspartates are shown in dark red in the mutant and salmon in the wild-type complex. Two principal conformational changes observed in the mutant structure compared to the WT are labeled 1 and 2. An enlarged view of conformational change 1, showing displacement of the NYN-1 loop containing R404 relative to RDE-8 is shown in the right panels. The mutant crystallized isomorphously with the wild-type complex, so these differences are not due to altered crystal packing. **(L)** Molecular dynamics simulations of the WT RDE-8/NYN-1 and RDE-8(S159R)/NYN-1 complexes. The distance between the $Mn^{2+}$ ion and each of the catalytic aspartates (D76, D157, D158, and D176) was monitored over time; the analysis for D157 is also shown in Figure 2G. Five independent simulations with a duration of 5 μs were performed for each complex. **(M)** Enlarged cartoon view of RDE-8(S159R)/NYN-1 catalytic site in a molecular dynamics simulation of the RDE-8(S159R)/NYN-1 complex (Rep2), just after D157 and D158 adopt the canonical active conformation (Time ~2.5 μs). Comparison with the RDE-8/NYN-1 crystal structure (this study, PDB: 28RS) is shown in the right panel, highlighting a similar conformational change to the one observed in (K).

#### Figure S3: Activation by MUT-15 uncovers RDE-8 cleavage preference

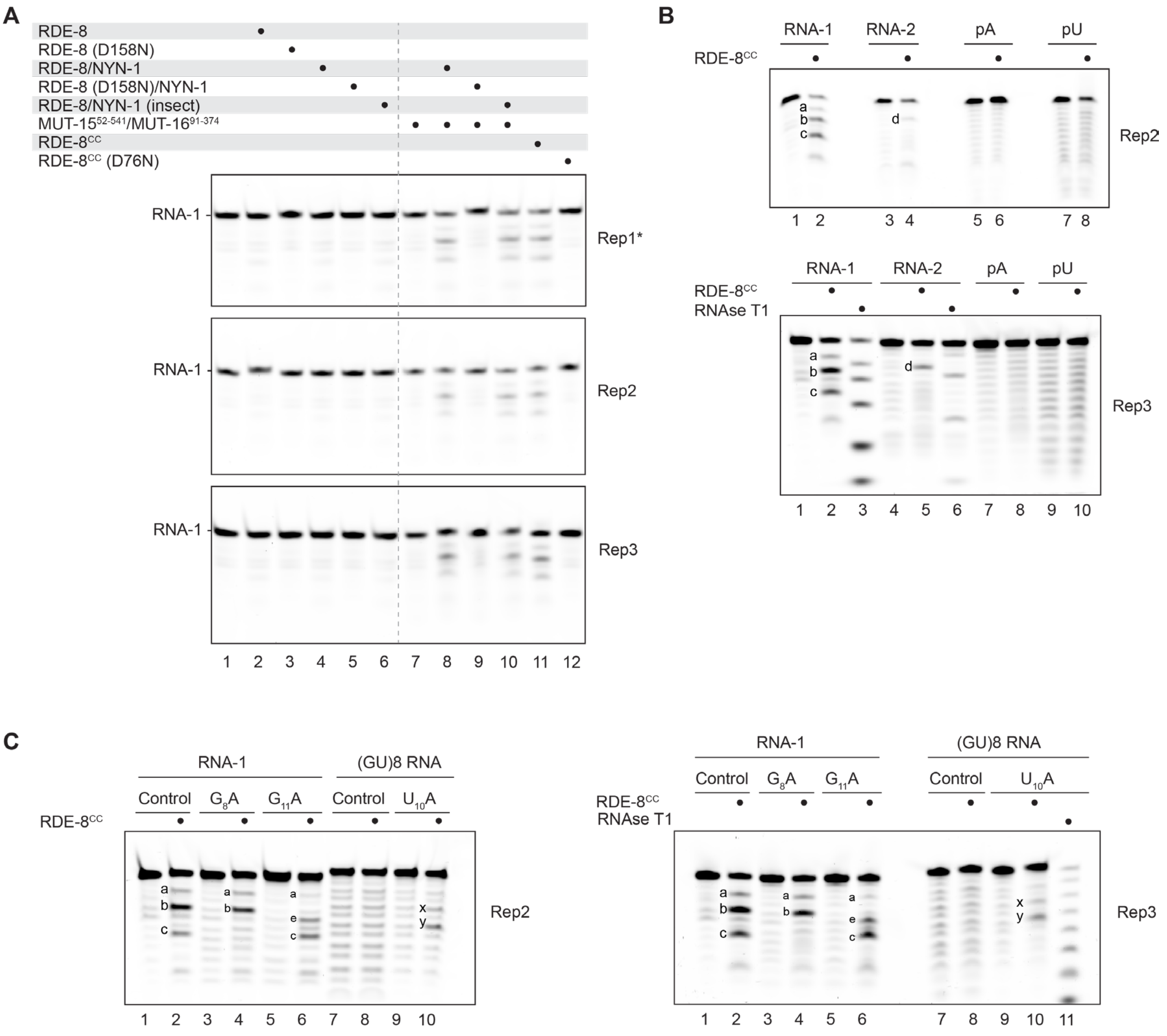

**(A–C)** Independent replicates of the RNA cleavage assays shown in Figure 3. Replicates 2 and 3 are indicated as Rep2 and Rep3; Rep1 corresponds to the experiment shown in the main figure. Cleavage fragments are marked as in Figure 3. **(A)** Replicates of the assays shown in Figure 3A and Figure S2J. The complete Rep1 gel is reproduced here for transparency; its right and left portions are also shown in Figure 3A and Figure S2J, respectively. **(B)** Replicates of the assays shown in Figures 3C and 3D. **(C)** Replicates of the assays shown in Figures 3E and 3F.

**Figure S4: MUT-15 remodels the nuclease module without relieving autoinhibition**

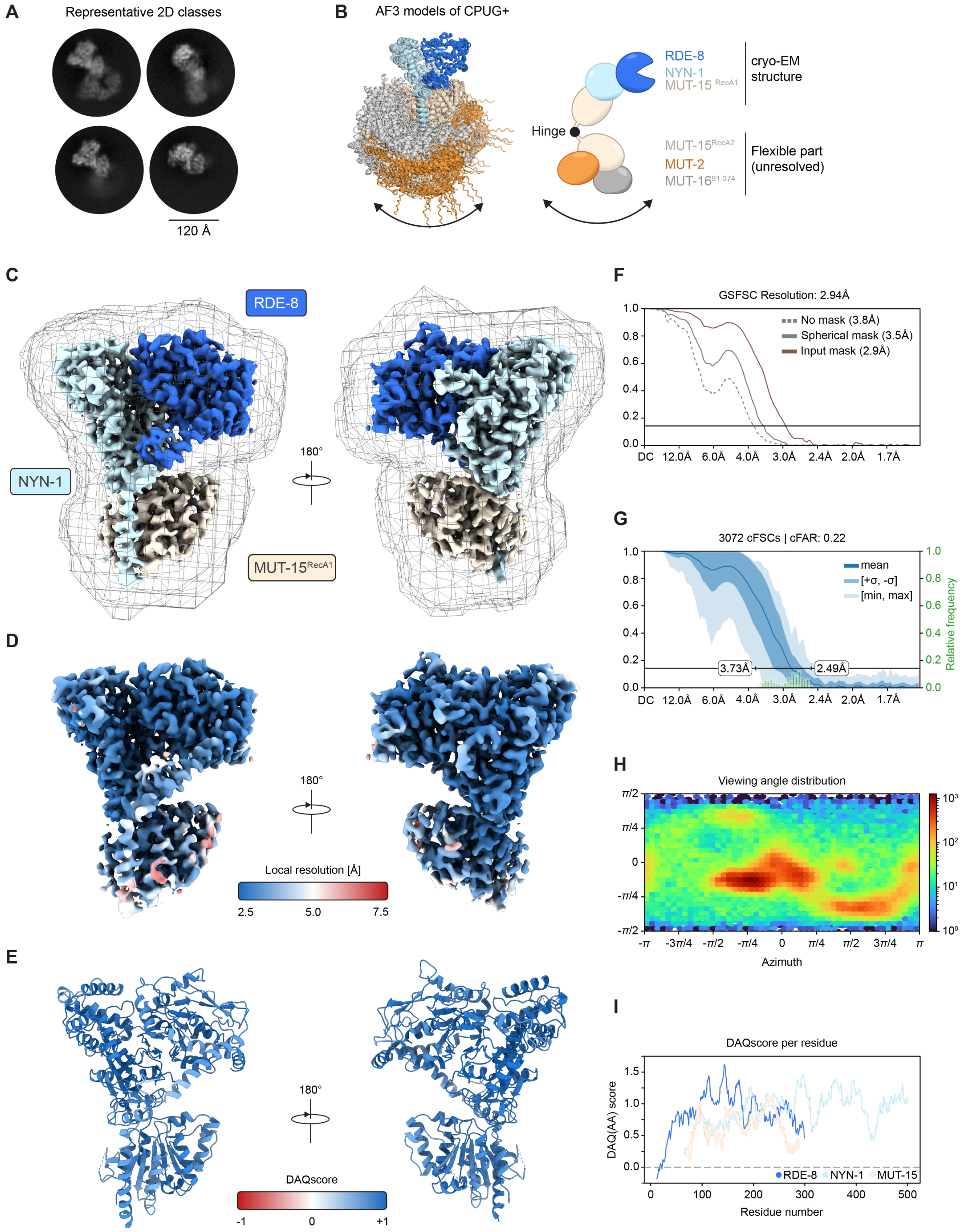

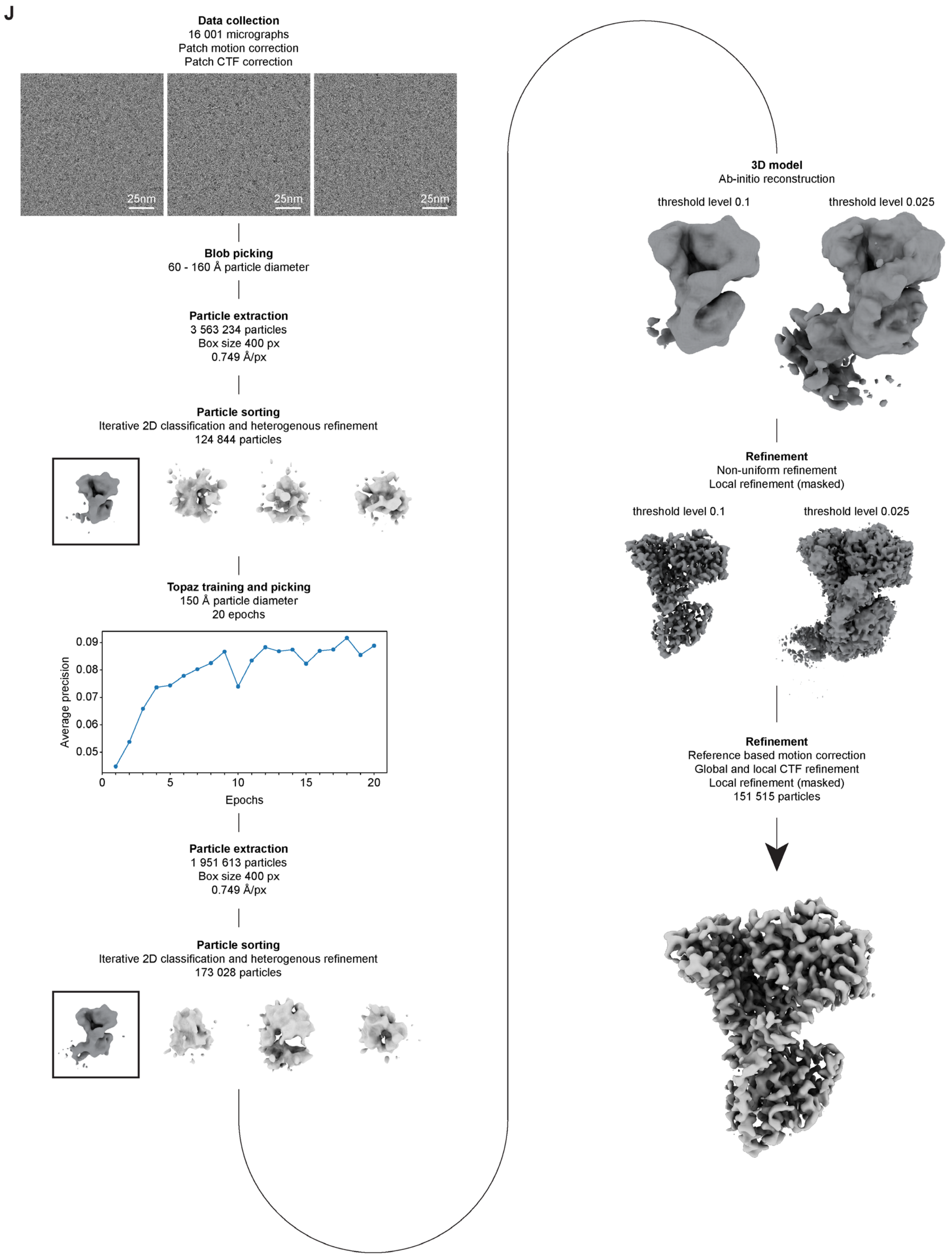
J
Data collection
16 001 micrographs
Patch motion correction
Patch CTF correction
25nm
25nm
25nm
Blob picking
60 - 160 Å particle diameter
Particle extraction
3 563 234 particles
Box size 400 px
0.749 Å/px
Particle sorting
Iterative 2D classification and heterogenous refinement
124 844 particles
Topaz training and picking
150 Å particle diameter
20 epochs
Average precision
0.09
0.08
0.07
0.06
0.05
0
5
10
15
20
Epochs
Particle extraction
1 951 613 particles
Box size 400 px
0.749 Å/px
Particle sorting
Iterative 2D classification and heterogenous refinement
173 028 particles
3D model
Ab-initio reconstruction
threshold level 0.1
threshold level 0.025
Refinement
Non-uniform refinement
Local refinement (masked)
threshold level 0.1
threshold level 0.025
Refinement
Reference based motion correction
Global and local CTF refinement
Local refinement (masked)
151 515 particles

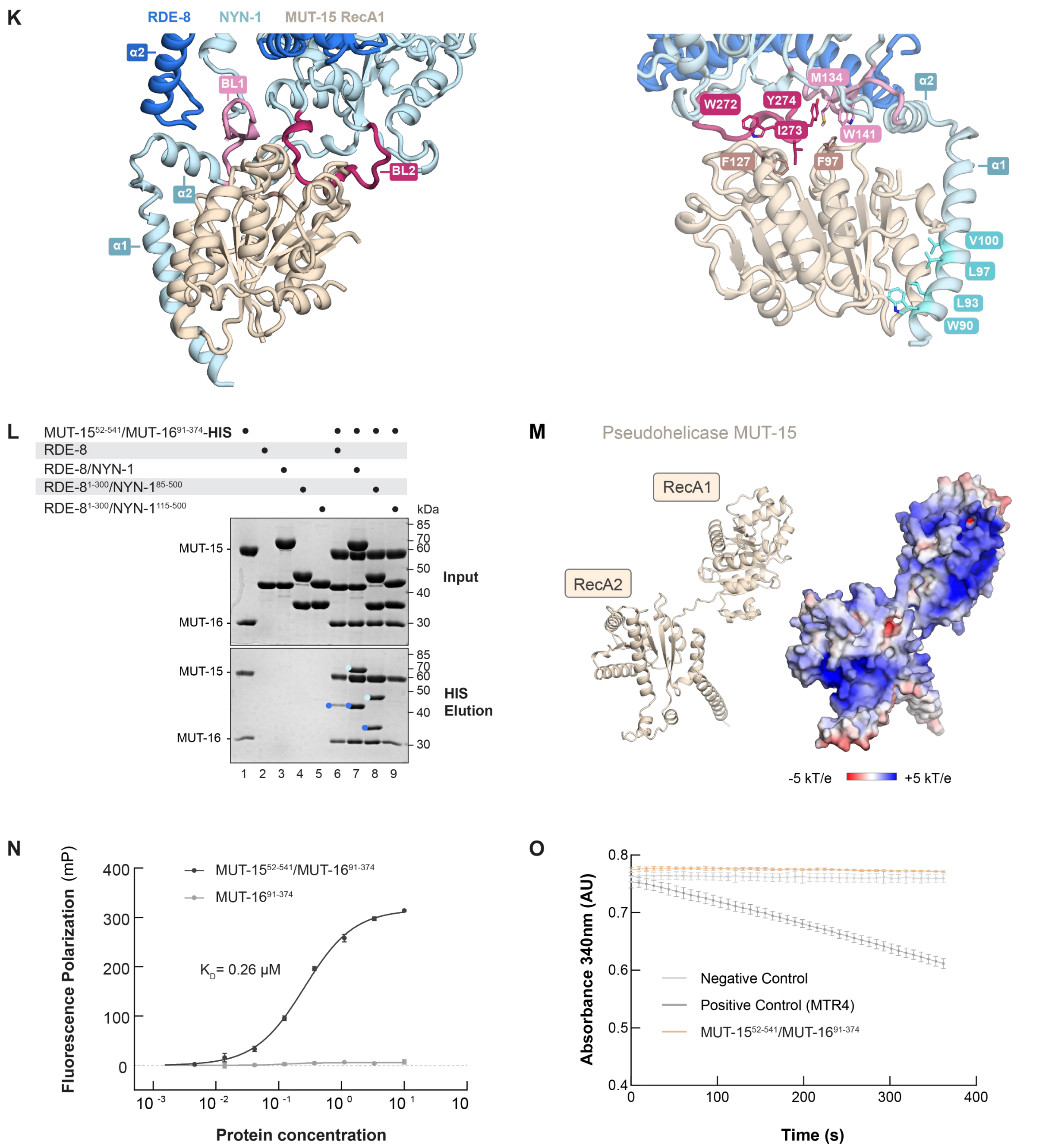

**(A)** Representative 2D classes used for cryo-EM reconstruction of the CPUG⁺ complex (PDB: 32VT). **(B)** Superposition of 25 AlphaFold3 predictions of CPUG⁺, highlighting flexibility of the lower region containing the MUT-15 RecA2-like domain, MUT-2, and MUT-16$^{91–374}$ relative to the upper region containing RDE-8, NYN-1, and the MUT-15 RecA1-like domain. The accompanying schematic illustrates the hinge between the two RecA-like domains of MUT-15 and the resulting flexibility that limited high-resolution reconstruction to the upper region of the complex. **(C)** Cryo-EM reconstruction of RDE-8/NYN-1/MUT-15$^{RecA1}$ heterotrimer at a global resolution of 2.9 Å colored by chain overlayed with the refinement mask (PDB: 32VT). **(D)** Local resolution estimation for the cryo-EM density of RDE-8/NYN-1/MUT-15 in the range of 2.5 to 7.5 Å. **(E)** Atomic model of RDE-8/NYN-1/MUT-15 colored by DAQ(AA) score. **(F)** Plot of the Gold Standard Fourier Shell Correlation (GSFSC) showing a global resolution estimation of 2.9 Å. **(G)** Plot of the conical Fourier Shell Correlation (cFSC) including the conical FSC Area Ratio (cFAR) of 0.14. **(H)** Plot of the viewing angle distribution. **(I)** Plot of the DAQ(AA) score per residue. **(J)** Processing workflow for the reconstruction of the RDE-8/NYN-1/MUT-15$^{RecA1}$ heterotrimer in cryoSPARC. **(K)** Interface between the RDE-8/NYN-1 nuclease module and MUT-15$^{RecA1}$ in the cryo-EM structure. Two enlarged views of the interface are shown; the right view highlights the interacting residues. NYN-1$^{α1}$ (89–111) binds within a hydrophobic groove on MUT-15$^{RecA1}$ through W90, L93, L97, and V100. Two aromatic-rich binding loops in the NYN-1 core, NYN-1$^{BL1}$ (128-143) and NYN-1$^{BL2}$ (271-278), form a second interface with MUT-15$^{RecA1}$. NYN-1 W141 in NYN-1$^{BL1}$ and W272, I273, and Y274 in NYN-1$^{BL2}$ pack against a hydrophobic surface on MUT-15$^{RecA1}$ centered on F97 and F127. NYN-1$^{α1}$, NYN-1$^{α2}$, NYN-1$^{BL1}$, and NYN-1$^{BL2}$ and RDE-8$^{α2}$ are highlighted. **(L)** Pull-down assay to test the interaction between the MUT-15$^{52–541}$/MUT-16$^{91–374}$ and RDE-8/NYN-1 complexes. His-tagged MUT-15$^{52–541}$ served as bait, and the indicated RDE-8/NYN-1 complexes were used as prey: full-length RDE-8/NYN-1, RDE-8$^{1–300}$/NYN-1$^{85–500}$, and RDE-8$^{1–300}$/NYN-1$^{115–500}$. RDE-8 alone was included as a control. Inputs and elutions were analyzed by SDS–PAGE and Coomassie staining. RDE-8 and NYN-1 constructs are indicated by dark- and light-blue dots, respectively. **(M)** Electrostatic potential of MUT-15 mapped on its AlphaFold3 prediction. The electrostatic surface potential was calculated by Adaptive Poisson–Boltzmann Solver from −5 kT/e (red) to +5 kT/e (blue). **(N)** RNA-binding affinities of MUT-15$^{52–541}$/MUT-16$^{91–374}$ determined by fluorescence polarization (FP) assays. MUT-16$^{91–374}$ is used as control. The 5′-FAM-labelled 16-ntRNA-1 (50 nM) was incubated with increasing concentrations of the indicated proteins. FP values were normalized by subtracting the FP values of the wells containing the labelled RNA alone. The mean of three experiments is shown, and error bars correspond to ± SD. The FP data are fitted to the Hill equation to obtain the dissociation constant, $K_d$. Best fit values: $K_d$=0.2559 μM, h=1.107, $B_{max}$=158 mP. **(O)** NADH-coupled ATPase assay of the MUT-15$^{52–541}$/MUT-16$^{91–374}$ complex (1μM). Decreased absorbance at 340 nm reflects NADH oxidation coupled to ATP hydrolysis. All reactions contained poly(U) RNA; the negative control lacked protein, and human MTR4 (residues 75–1042; 1μM) served as positive control. Data are shown as mean ± SD from five technical replicates.

#### Figure S5: RNA cleavage and tailing are coordinated within CPUG

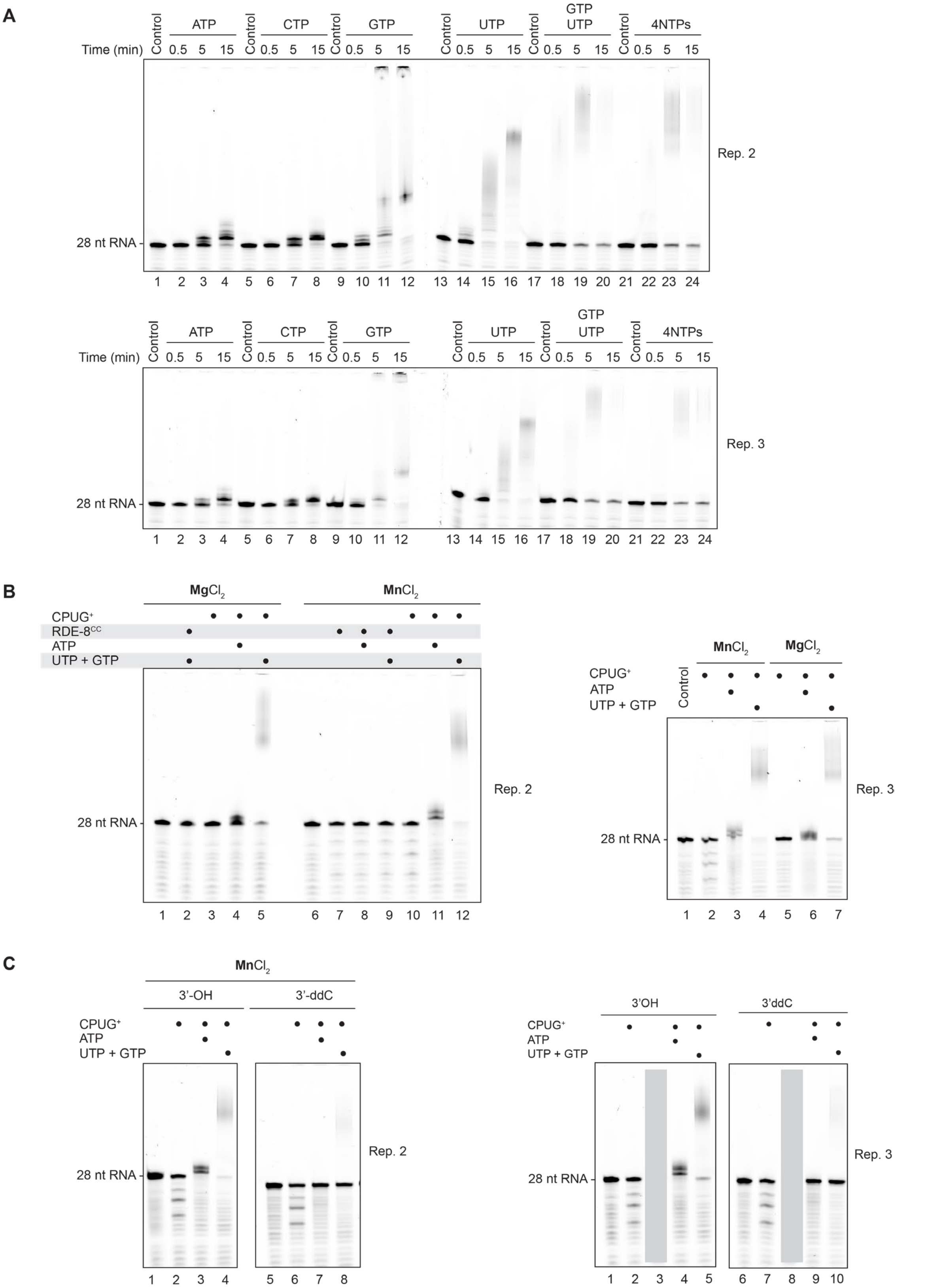

**(A-C)** Independent replicates of the assays shown in Figure 5. Replicates 2 and 3 are indicated as Rep2 and Rep3; Rep1 corresponds to the experiment shown in Figure 5. **(A)** Replicates of the assay shown in Figure 5A, including an additional time point at 15 min. **(B)** Replicates of the assay shown in Figure 5B. **(C)** Replicates of the assay shown in Figure 5C. The gray box covers lanes containing conditions not described in this study.

#### Figure S6: The substrate 3′ end determines the efficiency and register of poly(UG)ylation

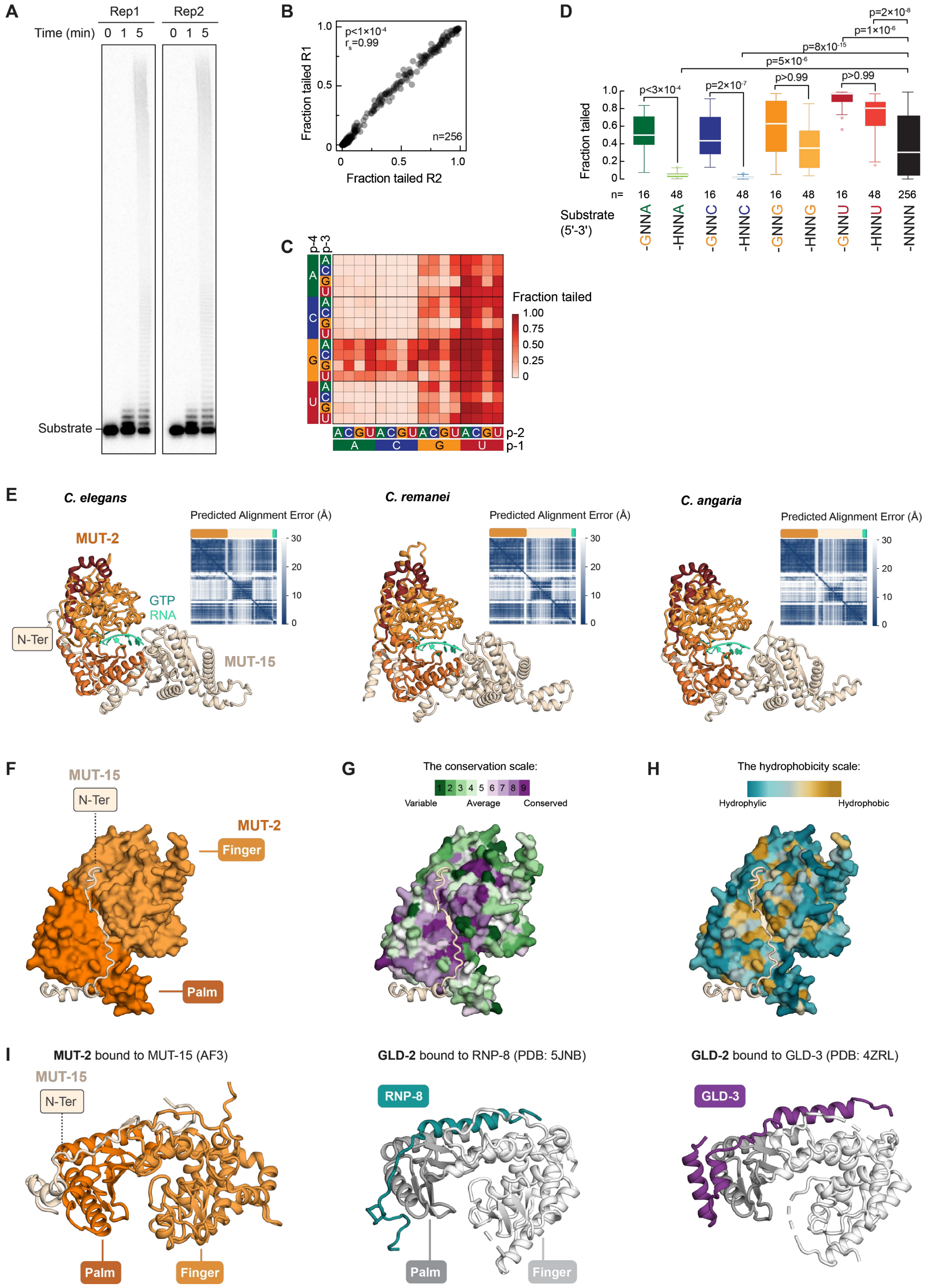

**(A)** RNA tailing assay using a 5′-radiolabeled 37-nt RNA library containing randomized 3′-terminal nucleotides (NNNN; 256 substrates), incubated with CPUG⁺. Reactions were performed with all four rNTPs at equimolar concentrations and sampled over time. Reaction products were resolved by denaturing polyacrylamide gel electrophoresis. The 5 min condition was chosen for sequencing experiments. The gel shows the two independent replicates used for sequencing and analysis. A zoom-in of Replicate 1 is also shown in Figure 6B. **(B)** Reproducibility of substrate tailing measurements between two independent replicates. Each point represents one substrate from the 4N RNA library. Tailing efficiency highly correlated between replicates (Spearman's $r_s = 0.99$, $p<1x10^{-4}$). **(C)** Heatmap showing the fraction tailed for all 256 substrates in the 4N library. Rows indicate nucleotide identity at position -1 (p-1) and position -2 (p-2) of the randomized sequence, whereas columns indicate nucleotide identity at position -3 (p-3) and position -4 (p-4). Color intensity represents the fraction of tailed substrate. **(D)** Tukey box plots showing the distribution of tailing efficiencies (fraction tailed) to specifically assess the contribution of position -4 (p-4) of the randomized sequence. Substrates were sorted by 3′-terminal nucleotide identity (p-1) and then separated according to whether p-4 contained G or a non-G nucleotide (H=A, C, or U). Adjusted p-values were calculated by Kruskal–Wallis test with Dunn's multiple-comparison correction. **(E)** AlphaFold3 predictions of the MUT-2/MUT-15 interaction in *C. elegans*, *C. remanei*, and *C. angaria* in the presence of a 5'-GUGU-3' RNA substrate, GTP, and two $Mg^{2+}$ ions. The predicted aligned error (PAE) plot for the top-ranked model is shown. For MUT-15, only the N-terminal region and RecA2-like domain are displayed. Protein identifiers: MUT-2, *C. elegans* (O44768), *C. remanei* (E3LUX7), and *C. angaria* (A0A9P1MVZ8); MUT-15, *C. elegans* (Q22061), *C. remanei* (E3NCK2), and *C. angaria* (A0A9P1N551). **(F-H)** Views of the interface between the MUT-15 N-terminal region and MUT-2 in the AlphaFold3 predicted *C. elegans* complex. **(F)** The palm and fingers domains of MUT-2 are highlighted, as in Deng et al. [6]. **(G)** The surface of MUT-2 is colored according to residue-conservation scores calculated using ConSurf. Multiple-sequence alignments of the following MUT-2 orthologs were used: *C. elegans* (O44768), *C. briggsae* (A8XG46), *C. remanei* (E3LUX7), *C. brenneri* (G0NU13), *C. japonica* (A0A8R1I1L6), *C. angaria* (A0A9P1MVZ8), and *C. auriculariae* (A0A8S1HDE2). **(H)** The surface of MUT-2 is colored according to hydrophobicity. (**I**) Comparison of the interface between the MUT-15 N-terminal region and MUT-2 from the AlphaFold3 prediction (left) with the crystal structures of GLD-2 bound to RNP-8 (PDB: 5JNB; middle)[7] or GLD-3 (PDB: 4ZRL; right)[8]. Palm and finger domains of the nucleotidyltransferases MUT-2 and GLD-2 are indicated.

### Supplemental Tables

#### Table S1: Recombinant proteins and purification strategies used in this study

H20 = 20 mM HEPES, pH 7.5; N150 = 150 mM NaCl; G10 = 10% glycerol; TCEP0.5 = 0.5 mM TCEP; DTT2 = 2 mM DTT.

| Name | Usage | Plasmid # | Protein | Residues | Mutations | N-Ter Tags | C-Ter Tags | Tag Cleaved | Method | Cells | Temp (°C) | Purification Steps | SEC Buffer |
|---|---|---|---|---|---|---|---|---|---|---|---|---|---|
| RDE-8 | pulldown | Ce30 | RDE-8 | FL | | HIS-GST-3C | | YES | | BL21 | 18 | HIS, REV HIS, Q, SEC | H20 N150 DTT2 |
| RDE-8 (D158N) | nuclease assays | Ce386 | RDE-8 | FL | D158N | HIS-GST-3C | | YES | | BL21 | 18 | HIS, REV HIS, Q, SEC | H20 N150 DTT2 |
| RDE-8/NYN-1 | pulldown | Ce30 | RDE-8 | FL | | HIS-GST-3C | | YES | co-transformatio | BL21 | 18 | HIS, REV HIS, Q, SEC | H20 N150 DTT2 |
| | | Ce31 | NYN-1 | FL | | HIS-TRX-3C | | YES | | | | | |
| RDE-8(D158N)/NYN-1 | nuclease assays | Ce386 | RDE-8 | FL | D158N | HIS-GST-3C | | YES | co-transformatio | LOBSTR | 18 | HIS, REV HIS, Q, SEC | H20 N150 DTT2 |
| | | Ce31 | NYN-1 | FL | | HIS-TRX-3C | | YES | | | | | |
| RDE-8(H181A)/NYN-1 | nuclease assays | Ce368 | RDE-8 | FL | H181A | HIS-GST-3C | | YES | co-transformatio | BL21 | 18 | HIS, REV HIS, Q, SEC | H20 N150 DTT2 |
| | | Ce31 | NYN-1 | FL | | HIS-TRX-3C | | YES | | | | | |
| RDE-8(S159R)/NYN-1 | nuclease assays | Ce369 | RDE-8 | FL | S159R | HIS-GST-3C | | YES | co-transformatio | BL21 | 18 | HIS, REV HIS, Q, SEC | H20 N150 DTT2 |
| | | Ce31 | NYN-1 | FL | | HIS-TRX-3C | | YES | | | | | |
| RDE-8/NYN-1(R404A) | nuclease assays | Ce30 | RDE-8 | FL | | HIS-GST-3C | | YES | co-transformatio | BL21 | 18 | HIS, REV HIS, Q, SEC | H20 N150 DTT2 |
| | | Ce372 | NYN-1 | FL | R404A | HIS-TRX-3C | | YES | | | | | |
| RDE-8$_{1-300}$/NYN-1$_{85-500}$ | pulldown | Ce371 | RDE-8 | 1-300 | D158N | HIS-MBP-3C | | | co-transformatio | BL21 | 18 | HIS, REV HIS, Hep, | H20 N150 DTT2 |
| | | Ce352 | NYN-1 | 85-500 | | HIS-MBP-3C | | | | | | | |
| RDE-8(S159R)/NYN-1 | crystal structure | Ce471 | RDE-8 | 1-300 | S159R | HIS-MBP-3C | | YES | co-transformatio | BL21 | 18 | HIS, REV HIS, HEP, | H20 N150 TCEP0.5 |
| | | Ce353 | NYN-1 | 115-500 | | HIS-MBP-3C | | YES | | | | | |
| RDE-8$_{1-300}$/NYN-1$_{115-500}$ | pulldown, crystal structure | Ce354 | RDE-8 | 1-300 | | HIS-MBP-3C | | YES | co-transformatio | BL21 | 18 | HIS, REV HIS, HEP, | H20 N150 TCEP0.5 |
| | | Ce353 | NYN-1 | 115-500 | | HIS-MBP-3C | | YES | | | | | |
| RDE-8/NYN-1 (insect) | nuclease assays | Ce431 | NYN-1 | FL | | | | YES | BigBac | Hi5 | 21 | Strep, 3C, Q, SEC | H20 N150 G10 DTT2 |
| | | | RDE-8 | FL | | Twin StrepII | | YES | | | | | |
| MUT-15$_{52-541}$/MUT16$_{91-374}$ | pulldown | Ce440 | MUT-16 | 91-374 | | GST-3C | | YES | BigBac | Hi5 | 21 | HIS, HEP, SEC | H20 N150 G10 DTT2 |
| | | | MUT-15 | 52-541 | | | HIS | NO | | | | | |
| RDE8$_{CC}$ | nuclease and tailing assays | Ce421 | MUT-16 | 91-374 | | GST | | YES | BigBac | Hi5 | 21 | Strep, 3C, Hep, SEC | H20 N150 G10 DTT2 |
| | | | MUT-15 | 52-541 | | | HIS | NO | | | | | |
| | | | NYN-1 | FL | | | | | | | | | |
| | | | RDE-8 | FL | | Twin StrepII | | YES | | | | | |
| RDE8$_{CC}$ (D76N) | nuclease assays | Ce439 | MUT-16 | 91-374 | | GST | | YES | BigBac | Hi5 | 21 | Strep, 3C, Hep, SEC | H20 N150 G10 DTT2 |
| | | | MUT-15 | 52-541 | | | HIS | NO | | | | | |
| | | | NYN-1 | FL | | | | | | | | | |
| | | | RDE-8 | FL | D76N | Twin StrepII | | YES | | | | | |
| CPUG$_{+}$ | nuclease and tailing assays | Ce433 | MUT-16 | 91-374 | | GST | | YES | BigBac | Hi5 | 21 | Strep, 3C, Hep, SEC | H20 N150 G10 DTT2 |
| | | | MUT-15 | FL | | | HIS | NO | | | | | |
| | | | NYN-1 | FL | | | | | | | | | |
| | | | RDE-8 | FL | | Twin StrepII | | YES | | | | | |
| | | | MUT2 | FL | | MBP | | YES | | | | | |

#### Table S2: Data collection and refinement statistics

| | **RDE-8/NYN-1** (PDB: 28RS) | **RDE-8(S159R)/NYN-1** (PDB: 28RV) |
|---|---|---|
| ***Data collection*** | | |
| Space group | $C222_1$ | $C222_1$ |
| Cell dimensions | | |
| a, b, c (Å) | 79.76, 103.05, 219.59 | 79.89, 103.69, 218.06 |
| α, β, γ (°) | 90, 90, 90 | 90, 90, 90 |
| Resolution (Å) | 39.24–1.65 (1.67–1.65) | 47.73–2.00 (2.03–2.00) |
| $R_{merge}$ | 0.157 (6.67) | 0.244 (1.15) |
| $R_{pim}$ | 0.044 (1.83) | 0.100 (0.468) |
| $CC_{1/2}$ | 0.998 (0.28) | 0.984 (0.69) |
| I / σI | 8.3 (0.4) | 4.8 (1.3) |
| Completeness (%) | 99.5 (98.3) | 99.7 (99.7) |
| Redundancy | 13.8 (14.0) | 6.9 (6.9) |
| ***Refinement*** | | |
| Resolution (Å) | 39.24–1.65 | 47.73–2.00 |
| No. reflections | 108,031 | 61,267 |
| $R_{work}$/ $R_{free}$ | 0.181 / 0.210 | 0.199 / 0.236 |
| No. atoms | 6,161 | 5,865 |
| Protein | 5,635 | 5,417 |
| Ligand/ion | 33 | 21 |
| Water | 493 | 427 |
| B-factors (Å²) | 36.1 | 28.8 |
| Protein | 35.7 | 28.6 |
| Ligand/ion | 47.2 | 34.9 |
| Water | 39.4 | 30.7 |
| R.m.s. deviations | | |
| Bond lengths (Å) | 0.007 | 0.002 |
| Bond angles (°) | 0.87 | 0.54 |
| Ramachandran (%) | | |
| Favored | 97.4 | 96.5 |
| Allowed | 2.4 | 3.2 |
| Outliers | 0.3 | 0.3 |
| Rotamer outliers (%) | 0.2 | 0.2 |
| Clashscore | 1.5 | 1.4 |

*Each structure was determined from a single crystal. Values in parentheses are for the highest-resolution shell.*

#### Table S3: Molecular dynamics simulations of wild-type RDE-8/NYN-1 and RDE-8(S159R)/NYN-1

Five independent replicates of 5 µs were performed per variant (50 µs aggregated).

| Simulation | Force field | Water | Box type | Solute-edge distance (nm) | Atoms | $Mn^{2+}$ | Counterions | Production (µs) |
|---|---|---|---|---|---|---|---|---|
| RDE-8 WT – NYN-1 1 | AMBER99SB-ILDN | TIP3P | octahedron | 1.5 | 100922 | 1 | 2 $Cl^-$ | 5 |
| RDE-8 WT – NYN-1 2 | AMBER99SB-ILDN | TIP3P | octahedron | 1.5 | 100922 | 1 | 2 $Cl^-$ | 5 |
| RDE-8 WT – NYN-1 3 | AMBER99SB-ILDN | TIP3P | octahedron | 1.5 | 100922 | 1 | 2 $Cl^-$ | 5 |
| RDE-8 WT – NYN-1 4 | AMBER99SB-ILDN | TIP3P | octahedron | 1.5 | 100922 | 1 | 2 $Cl^-$ | 5 |
| RDE-8 WT – NYN-1 5 | AMBER99SB-ILDN | TIP3P | octahedron | 1.5 | 100922 | 1 | 2 $Cl^-$ | 5 |
| RDE-8 S159R – NYN-1 1 | AMBER99SB-ILDN | TIP3P | octahedron | 1.5 | 100921 | 1 | 3 $Cl^-$ | 5 |
| RDE-8 S159R – NYN-1 2 | AMBER99SB-ILDN | TIP3P | octahedron | 1.5 | 100921 | 1 | 3 $Cl^-$ | 5 |
| RDE-8 S159R – NYN-1 3 | AMBER99SB-ILDN | TIP3P | octahedron | 1.5 | 100921 | 1 | 3 $Cl^-$ | 5 |
| RDE-8 S159R – NYN-1 4 | AMBER99SB-ILDN | TIP3P | octahedron | 1.5 | 100921 | 1 | 3 $Cl^-$ | 5 |
| RDE-8 S159R – NYN-1 5 | AMBER99SB-ILDN | TIP3P | octahedron | 1.5 | 100921 | 1 | 3 $Cl^-$ | 5 |

#### Table S4: Cryo-EM data collection, processing and validation statistics

| | **RDE-8/NYN-1/MUT-15** *(EMDB-59208, PDB 32VT)* |
|---|---|
| ***Data collection and processing*** | |
| Microscope | Thermo Fisher Titan Krios G4 |
| Voltage (kV) | 300 |
| Detector | Thermo Fisher Falcon 4i |
| Magnification | 165,000 |
| Pixel size (Å) | 0.749 |
| Electron dose ($e^{-}$/Å$^2$) | 40 |
| EER fractions (#) | 40 |
| Defocus range (µm) | −2.5 to −1.0 |
| Energy filter slit width (eV) | 10 |
| Micrographs collected (#) | 16,001 |
| Initial particle images (#) | 3,563,234 |
| Final particle images (#) | 151,515 |
| Box size (px) | 400 × 400 |
| Symmetry imposed | C1 |
| Map sharpening B factor (Å$^2$) | n/a |
| Map resolution (Å) | 2.94 |
| FSC threshold | 0.143 |
| Map resolution range, 95th / 5th percentile (Å) | 2.77–8.71 |
| FSC threshold | 0.5 |
| ***Refinement*** | |
| Initial model used | AlphaFold3 |
| Software used | Phenix, Coot, ISOLDE |
| ***Model composition*** | |
| Chains (#) | 3 |
| Non-hydrogen atoms (#) | 7,342 |
| Protein residues (#) | 904 |
| Nucleic acid bases (#) | 0 |
| Ligands (#) | 0 |
| Water (#) | 0 |
| ***B factors (Å$^2$)*** | |
| Protein min / max / mean | 47.93 / 190.53 / 98.09 |
| ***R.m.s. deviations*** | |
| Bond lengths (Å) | 0.003 |
| Bond angles (°) | 0.672 |
| ***Validation*** | |
| MolProbity score | 1.49 |
| Clashscore | 4.96 |
| Rotamer outliers (%) | 0.97 |
| ***Ramachandran*** | |
| Favored (%) | 96.52 |
| Allowed (%) | 3.25 |
| Outliers (%) | 0.22 |

##### Table S5: Processed sequencing data from CPUG RNA tailing assays.

Processed sequencing data from two biological replicates of *in vitro* tailing experiments. Related to Figure 6 and Figure S6. The Excel workbook contains four worksheets. Data_All2Reps_q20norm contains the sequencing results for each of the 256 possible 4N substrate sequences at 0 and 5 min in both replicates. For each substrate, the sheet reports sequencing read counts and RPM-normalized read counts, the fraction of untailed and tailed reads, and RPM values for reads carrying tails of 1–10 nt. SummedTailCnt5_q20 summarizes, for the 5-min samples, the counts and relative frequencies of tails initiating with A, C, G, or U for each substrate and replicate. Replicate1_tail content and Replicate2_tail content list the individual tail sequences detected for each 4N substrate, together with substrate read counts, tail read counts, tail length, and RPM-normalized read counts- for both substrate and tails.

##### Movie S1. MUT-15 binding is associated with remodeling of the nuclease module

The movie shows an interpolated transition between the crystal structure of the RDE-8/NYN-1 complex (PDB: 28RS) and the cryo-EM structure of the RDE-8/NYN-1/MUT-15$^{RecA1}$ complex (PDB: 32VT). Proteins are colored as in Figure 4. MUT-15 is shown approaching from a displaced position while RDE-8/NYN-1 transitions from the crystal to the cryo-EM conformation. The trajectory of MUT-15 is illustrative and does not represent an experimentally determined binding pathway. The NYN-1 helix resolved in the cryo-EM structure but absent from the crystal structure appears upon formation of the final complex.

### Supplemental Material and Methods

##### Pull-Down Assays

Protein–protein interactions were analyzed by Ni–NTA pull-down assays using purified proteins in incubation buffer containing 20 mM HEPES/NaOH, 150 mM NaCl, 10% (v/v) glycerol, 20 mM imidazole, 2 mM DTT, and 0.05% NP-40. MUT-15$^{52–541}$/MUT-16$^{91–374}$ was used at a final concentration of 4 µM and prey proteins at 7.5 µM. Bait and prey proteins were mixed and incubated for 30 min on ice before addition to 15 µl Ni–NTA agarose beads in a final reaction volume of 80 µl. Samples were incubated for 2 h at 4°C with gentle agitation. Beads were washed once with incubation buffer and three times with wash buffer containing 20 mM HEPES/NaOH, 250 mM NaCl, 10% (v/v) glycerol, 20 mM imidazole, 2 mM DTT, and 0.05% NP-40. Bound proteins were eluted in 50 µl Ni–NTA elution buffer containing 50 mM $NaH_2PO_4$ (pH 8.0), 25 mM Tris-HCl (pH 7.5), 250 mM NaCl, 500 mM imidazole, 10% glycerol, and 5 mM β-mercaptoethanol by incubation for 10 min. Input and elution samples were mixed with Laemmli loading dye and analyzed by SDS–PAGE on 12% polyacrylamide gels. Input lanes contained 12.5% of the total input material, and elution lanes contained 13.3% of the total elution volume.

##### ATPase assays

ATPase activity of the MUT-15$^{52–541}$/MUT-16$^{91–374}$ complex was measured using an NADH-coupled assay in 384-well plates, on a Spark multimode microplate reader (Tecan). Reactions were performed in a final volume of 60 µL containing 50 mM HEPES pH 7.5, 50 mM potassium acetate, 5 mM $MgCl_2$, 2 mM DTT, 0.1 mg/mL BSA, 25 µg poly(U) RNA, 3 mM phosphoenolpyruvate, 1:40 dilution of pyruvate kinase/lactate dehydrogenase suspension, 400 µM NADH, and 1 µM protein. Reaction mixture lacking ATP were incubated for 20 min at room temperature (22°C) followed by 5 min at 30°C in the plate reader; reactions were then started by addition of 1 mM ATP. ATP hydrolysis was monitored by following the decrease in absorbance at 340 nm resulting from NADH oxidation at 30°C. Reactions containing RNA but no protein served as negative controls, whereas human MTR4$^{(75–1042)}$ was used as a positive control. Five replicate reactions were measured per condition in parallel on the same plate.

#### Fluorescence Polarization

Fluorescence Polarization experiments were performed as described in Busetto et al. [9]. A 16-nt single-stranded RNA-1 (5'-GUUGAUUGAAGAGUUC-3') carrying a 5′-6-fluorescein amidite (5′-FAM) label was purchased from Ella Biotech (Fuerstenfeldbruck, Germany). Increasing concentrations of protein were incubated with 50 nM RNA in a volume of 20 µl at room temperature for 30 min. The buffer contained 20 mM HEPES/NaOH pH 7.5, 150 mM NaCl, 10 % [v/v] glycerol and 5 mM EDTA. Fluorescence polarization data were recorded on a Tecan SPARK plate reader using an excitation wavelength of 485 nm with a bandwidth of 20 nm and an emission wavelength of 535 nm with a bandwidth of 25 nm. Millipolarization (mP) values were normalized by subtracting the mP values of wells containing only the fluorophore-labelled RNA. Data were analyzed by nonlinear regression using the Hill equation in GraphPad Prism version 10. The mean of three replicates ± SD is shown.

#### Mass spectrometry characterization of CPUG$^+$

##### Intact protein liquid chromatography-mass spectrometry

LC-MS analysis was performed on a Vanquish™ Horizon UHPLC system (Thermo Scientific) coupled to a Synapt G2-Si mass spectrometer (Waters) equipped with a ZSpray ESI source (Waters). 230 ng of protein in 3 µL solution was loaded to an XBridge Protein BEH C4 column (300Å, 2.5 µm particle size, dimensions 2.1 mm x 150 mm; Waters) with a working temperature of 50°C, 0.1% formic acid (FA) as solvent A, 100% acetonitrile, 0.08% FA as solvent B. Proteins were separated on a 6 min step gradient from 12-40-64 % solvent B at a flow rate of 250 µL/min. The mass spectrometer was operated in resolution mode, the capillary voltage was set to 2 kV, the source temperature was 120°C, the desolvation temperature 400°C. Data were recorded with MassLynx V 4.2 (Waters) and calibrated by the lockmass of Glu-Fibrinopeptide B. Raw data were analyzed using the MaxEnt 1 process to reconstruct the uncharged average protein mass.

##### Bottom-up liquid chromatography-mass spectrometry

Protein solution was diluted with an 8 M urea in 50 mM ABC stock solution to a final urea concentration of 6 M. 10 mM DTT was added followed by incubation for 30 min at room temperature. Reduced thiols were alkylated by adding iodoacetamide (500 mM stock solution in dH2O) and let react for 30 min at room temperature in the dark. Residual iodoacetamide was quenched with half the amount of DTT used in the reduction step. Prior to adding proteases, the solution was diluted to 1 M urea with 50 mM ABC. Proteases were added at a 1:30 (protease:protein) ratio and incubated at 37°C overnight (for trypsin) or at 25°C for 2-5 h for chymotrypsin. Digestion was stopped by acidifying the solution with 10% TFA. Peptides were purified on StageTips according to Rappsilber et al. [10] with some modifications. Three C18 Empore disk punches were packed into 200 µL pipette tips. All applied solutions were passed through the tip by centrifugation. Tips were prewetted with 100 µL methanol, followed by 100 µL 80% acetonitrile (ACN), 0.1% TFA, and equilibrated with 100 µL 0.1% TFA by spinning at 376 g for 4-6 min. Acidified peptide solutions were applied, and spun at 271 g till passed through. Bound peptides were washed with 100 µL 0.1% TFA, and eluted with 2 x 30 µL 40% ACN, 0.1% TFA into PCR tubes. Eluates were reduced to dryness in a vacuum centrifuge and taken up in 20 µL 2% ACN, 0.1% TFA.

LC-MS setup was as for the cross-link samples, except for measuring without FAIMS and running a 60 min gradient. The mass spectrometer was operated in data-dependent acquisition (DDA) mode with 2 s cycle time. Survey scans were acquired from 375-2000 m/z, with normalized AGC target of 300%, resolution of 120,000. The most intense precursor ions (charge states +2 to +6) were selected for fragmentation using an isolation window of 1.4 m/z. Selected ions were analyzed with a maximum fill time of 100 ms, normalized AGC target of 200%, and resolution of 30,000 after HCD fragmentation with a collision energy of 30%. Monoisotopic precursor selection (MIPS) was set to "peptide" mode, the intensity threshold to 2.5E4, and selected precursors were dynamically excluded for 20 s with isotope exclusion enabled.

MS raw data were analyzed with FragPipe running the standard workflow, except “Normalize intensity across runs” and “Match between runs” were turned off, and maximal peptide size was extended to 80 AA respect 8000 Da. Cleavage specificity was set to trypsin with 2 missed cleavages allowed. The FDR was set to 1%. Oxidation of methionine, phosphorylation on serine, threonine and tyrosine and N-terminal protein acetylation were specified as variable modifications. Carbamidomethylation of cysteine was set as a fixed modification. MS2 spectra were searched against target sequences in their expression background (2024_01_7111_Trichoplusia_ni_21155entries.fasta), concatenated with a database of common laboratory contaminants (https://github.com/maxperutzlabs-ms/perutz-ms-contaminants ).

**Crosslinking Mass Spectrometry**

Protein complexes (5 µM) were cross-linked with 0.5 mM DSBU for 30 min at room temperature in a final volume of 25 µl. Reactions were quenched by addition of 50 mM TRIS/HCl pH 7.5. Half of the cross-linked sample was subjected to size-exclusion chromatography on a Superdex 200 Increase 3.2/300 column using a ÄKTAmicro system. Peak fractions corresponding to the cross-linked complex were analyzed by SDS–PAGE on an 8% Tris/Glycine gel. The corresponding band was excised from the gel and subjected to downstream analysis.

The Coomassie-stained gel band was destained with a mixture of ACN and 50 mM ammonium bicarbonate (ABC). The proteins were reduced using 10 mM DTT and alkylated with 50 mM iodoacetamide. Trypsin (Promega; Trypsin Gold, Mass Spectrometry Grade) was used for proteolytic cleavage. Digestion was carried out at 37°C overnight. Formic acid (FA, 10%) was used to stop digestion and 5% FA, 50% ACN was used to extract peptides from the gel.

LC-MS analysis was performed on a Vanquish Neo UHPLC system (Thermo Scientific) coupled to an Orbitrap Exploris 480 mass spectrometer (Thermo Scientific). The system was equipped with a FAIMS Pro (Thermo Scientific), a Nanospray Flex ion source (Thermo Scientific) with coated emitter tips (PepSep, MSWil), and a Butterfly Portfolio Heater (Phoenix S&T). Peptides were loaded onto a trap column (PepMap Neo C18 5 mm × 300 µm, 5 µm particle size, Thermo Scientific) using 0.1% trifluoroacetic acid (TFA) as mobile phase, and separated on an analytical column (Acclaim PepMap 100 C18 HPLC Column, 50 cm × 75 µm, 2 µm particle size, Thermo Scientific) applying a linear gradient starting with a mobile phase of 98% solvent A (0.1% FA) and 2% solvent B (80% ACN, 0.08% FA), increasing to 35% solvent B over 120 min at a flow rate of 230 nl/min. The analytical column was heated to 30°C.

The mass spectrometer was operated in DDA mode with FAIMS compensation voltage (CV) set to alternate between -40, -55 and -70 V, with 1 s cycle time per CV. Survey scans were acquired from 375-1600 m/z, with normalized AGC target of 100%, resolution of 60,000. The most intense precursor ions (charge states +3 to +8) were selected for fragmentation using an isolation window of 1.4 m/z. Selected ions were analyzed with a maximum fill time of 200 ms, normalized AGC target of 100%, and resolution of 30,000 after HCD fragmentation with stepped collision energy of 26%, 28%, 30%. Monoisotopic precursor selection (MIPS) was set to “peptide” mode, the intensity threshold to 2.5E4, and selected precursors were dynamically excluded for 45 seconds with isotope exclusion enabled.

The RAW MS data were analyzed with FragPipe (22.0), using MSFragger (4.3)[11], IonQuant (1.11.11)[12], and Philosopher (5.1.2)[13]. The default FragPipe workflow for label free quantification (LFQ-MBR) was used, except “Normalize intensity across runs” was turned off. Cleavage specificity was set to Trypsin/P, with two missed cleavages allowed. The protein FDR was set to 1%. A mass of 57.02146 (carbamidomethyl) was used as fixed cysteine modification; methionine oxidation, and protein N-terminal acetylation were specified as variable modifications. MS2 spectra were searched against the E. coli reference proteome from Uniprot (release 2025_01(ID25313), 2026_01(ID26031)), concatenated with a database of common laboratory contaminants (release 2025_01, https://github.com/maxperutzlabs-ms/perutz-ms-contaminants) and additional protein sequences corresponding to target constructs. To identify cross-linked peptides, the raw files were converted into mgf format using MSConvert (https://proteowizard.sourceforge.io/) and searched against the target sequences and 3 top non-contaminant proteins from the FragPipe search sorted by spectral counts using pLink software (3.0.16)[14]. DSBU was selected as the cross-linking chemistry. Carbamidomethyl on Cys was set as fixed, oxidation of Met, and protein N-terminal acetylation as variable modifications. Enzyme specificity was set to trypsin with 4 missed cleavages. Search results were filtered for 1% separate FDR at peptide pairs level, and a maximum e-value of 0.001. Cross-link maps were generated in Xinet (http://crosslinkviewer.org/).
